# Sniper2L, a high-fidelity Cas9 variant with high activity

**DOI:** 10.1101/2022.12.05.519240

**Authors:** Young-hoon Kim, Nahye Kim, Ikenna Okafor, Sungchul Choi, Seonwoo Min, Joonsun Lee, Keunwoo Choi, Janice Choi, Vinayak Harihar, Youngho Kim, Jin-Soo Kim, Jungjoon K. Lee, Taekjip Ha, Hyongbum Henry Kim

## Abstract

Although several high-fidelity SpCas9 variants that have reduced activities at mismatched target sequences have been reported, it has been observed that this increased specificity is associated with reduced on-target activity, limiting the applications of the high-fidelity variants when efficient genome editing is required. Here, we developed an improved version of Sniper-Cas9, Sniper2L, which represents an exception to this trade-off trend as it showed higher specificity with retained high activity. We evaluated Sniper2L activities at a large number of target sequences, and developed DeepSniper, a deep-learning model that can predict the activity of Sniper2L. We also confirmed that Sniper2L can induce highly efficient and specific editing at a large number of target sequences when it is delivered as a ribonucleoprotein complex. Mechanically, the high specificity of Sniper2L originates from its superior ability to avoid unwinding a target DNA containing even a single mismatch. We envision that Sniper2L will be useful when efficient and specific genome editing is required.

## Introduction

Applications of SpCas9-induced genome editing are often restricted due to off-target effects or insufficient on-target editing. Several high-fidelity variants, such as eSpCas9(1.1)^1^, Cas9-HF1^2^, HypaCas9^3^, Cas9_R63A/Q768A^4^, evoCas9^5^, HiFi Cas9^6^, and Sniper-Cas9^7^, have been developed. However, the modifications introduced in these variants to decrease off-target cleavage also hamper their general on-target cleavage activities, such that a trade-off between the general activity and specificity^8^ is observed when the variants are tested with a large number of target sequences. A high-fidelity variant that exhibits a general activity level similar to that of SpCas9 would facilitate applications of Cas9-based genome editing in areas including gene therapy and genetic screening.

In this study, we developed Sniper2L, a next-generation high-fidelity variant, using directed evolution of Sniper-Cas9. To evaluate the specificity and activity of Sniper2L at a large number of target sequences, we delivered it together with guide RNA using two different methods; lentiviral expression and electroporation of ribonucleoprotein (RNP) complex, a therapeutically relevant method. Our high-throughput evaluations showed that Sniper2L has higher fidelity than Sniper-Cas9 while retaining its general level of activity similar to the level of SpCas9, overcoming the trade-off between activity and specificity regardless of the delivery method. We believe that Sniper2L will facilitate applications of genome editing due to its high general activity and low levels of off-target effects.

## Results

### Directed evolution of Sniper-Cas9

Previously, we used “Sniper-screen” for directed evolution of Cas9 in *E. coli*^7^ (Supplementary Fig. 1). In brief, both positive [Cas9-mediated cleavage of a plasmid containing a lethal gene (*ccdB*)] and negative (lack of *E. coli*-killing cleavage at a mismatched off-target genomic site) selection pressure were applied to SpCas9 mutant libraries in which the entire Cas9-encoding sequence contained random errors (library complexity, up to 10^7^); a fragment of the human *EMX1* gene was used for the matched and mismatched target sequences. The initial Sniper-screen resulted in the identification of three Cas9 variants named Clone-1, Clone-2, and Clone-3^7^. We selected Clone-1 because it induced high frequencies of on-target indels with many different single-guide RNAs (sgRNAs) compared to Clone-2 and Clone-3, which showed low on-target indel efficiencies with the same sgRNAs. High indel frequencies were observed when these variants were tested with the sgRNA EMX1.3, which was used in the Sniper-screen. To distinguish Cas9 variants with reduced on-target activities such as Clone-2 and Clone-3 from those with maintained on-target activities, we needed to perform the Sniper-screen with a sgRNA that would result in low on-target indel efficiencies with Clone-2 and Clone-3 while retaining WT-level indel efficiencies with Clone-1. When we used EMX1.6 sgRNA, which was previously used to determine the specificity of Cas9^9^, we found that on-target activities of Clone-2 and -3 were dramatically decreased as compared to that of Clone-1 (Supplementary Figure 2). Thus, we chose EMX1.6 sgRNA for screening in the current study. As a mismatched target sequence, we used the target sequence with a mismatch in the 13^th^ position (gcgccacTggttgatgtgat; the mismatched nucleotide at position 13 is capitalized).

Libraries encoding mutant versions of Sniper-Cas9 with random errors in the Sniper-Cas9 sequence were constructed using the three different mutagenesis kits that were used in the previous Sniper-screen^7^. The Sniper-screen selection procedure was repeated four times with the EMX1.6 sgRNA (Supplementary Figure 3). The final clones were sequenced and a hot spot at the 1007^th^ amino acid of Sniper-Cas9 was identified (Supplementary Figure 4). We introduced all possible amino acid mutations at the 1007^th^ amino acid and measured the activities of these 19 mutants at matched and mismatched target sequences using another sgRNA (not EMX1.6) (Supplementary Fig. 5 and Supplementary Table 1). Among the 19 mutants, we selected six mutants that showed high on-target activity and low activity at the mismatched target sequence. We evaluated these six mutants at four different target sequences (Supplementary Fig. 6 and Supplementary Table 1). We finally selected two mutants named Sniper2L and Sniper2P with E1007L and E1007P mutations, respectively, both of which retained high on-target activities with diminished off-target activities at the tested four target sequences.

### High-throughput assessments of the activities and specificities of the Sniper2 variants

Although we compared the activities of Sniper2L and Sniper2P at four target sequences, to compare the general activities of these two variants, much more target sequences should be used^8^. To evaluate the activities of these two variants at a large number of target sequences, we adopted a high-throughput evaluation approach that we previously used to compare activities of various Cas9 variants^8^ (Supplementary Fig. 7a). For this high-throughput evaluations, we first generated individual cell lines, each containing a single copy of a variant-expressing lentivirus^8^, which led to comparable expression levels of Sniper1 and the Sniper2 variants (Supplementary Fig. 7b). We then transduced our previously-described lentiviral libraries of pairs of sgRNA-encoding and corresponding target sequences^8, 10^, into the Sniper-Cas9 variant-expressing cells and indel frequencies at the integrated target sequences were determined by deep sequencing four and seven days after the transduction of lentiviral libraries (Methods). The used libraries were libraries A, B, and C^8^, which contained 11,802, 23,679, and 7,567 sgRNA-target pairs, respectively. In brief, library A included 8,130 and 3,672 pairs to evaluate protospacer adjacent motif (PAM) compatibility and mismatch tolerance, respectively (Supplementary Table 2). Library B, which contained 8,744, 12,093, and 2,660 pairs with NGG, NGH, and non-NG PAMs, respectively, was used for validating variants at a large number of target sequences (Supplementary Table 3). In contrast to libraries A and B, Library C utilized perfectly matched N_20_ sgRNAs generated by transfer RNA (tRNA)-associated processing (hereafter, tRNA-N_20_ sgRNAs), with the majority of target sequences taken from library B (Supplementary Table 4). Because indel frequencies between two technical replicates were well correlated (Supplementary Fig. 8), we combined the read counts from two replicates to draw more accurate conclusions^8^.

We first determined the PAM compatibilities recognized by the Sniper2 variants using library A containing target sequences with NNNN PAMs. We found that the PAM compatibilities of Sniper-Cas9 variants were identical and that the highest average activities were observed at target sequences with NGG PAMs (Supplementary Fig. 9). These results are in line with the PAM compatibilities of other high-fidelity variants^8^ and would be attributable to the lack of mutations within the PAM-interacting domain of Sniper-Cas9 variants. Based on these results, target sequences with NGG PAMs were chosen for subsequent analysis.

We evaluated the activities of the Sniper2 variants at a large number of matched and mismatched target sequences. For assessing on-target activities, the 8,744 target sequences with NGG PAMs in library B were utilized. We found that Sniper2L exhibited significantly higher efficiencies than Sniper1, whereas Sniper2P induced the lowest indel frequencies (Fig. 1a).

**Figure 1.**
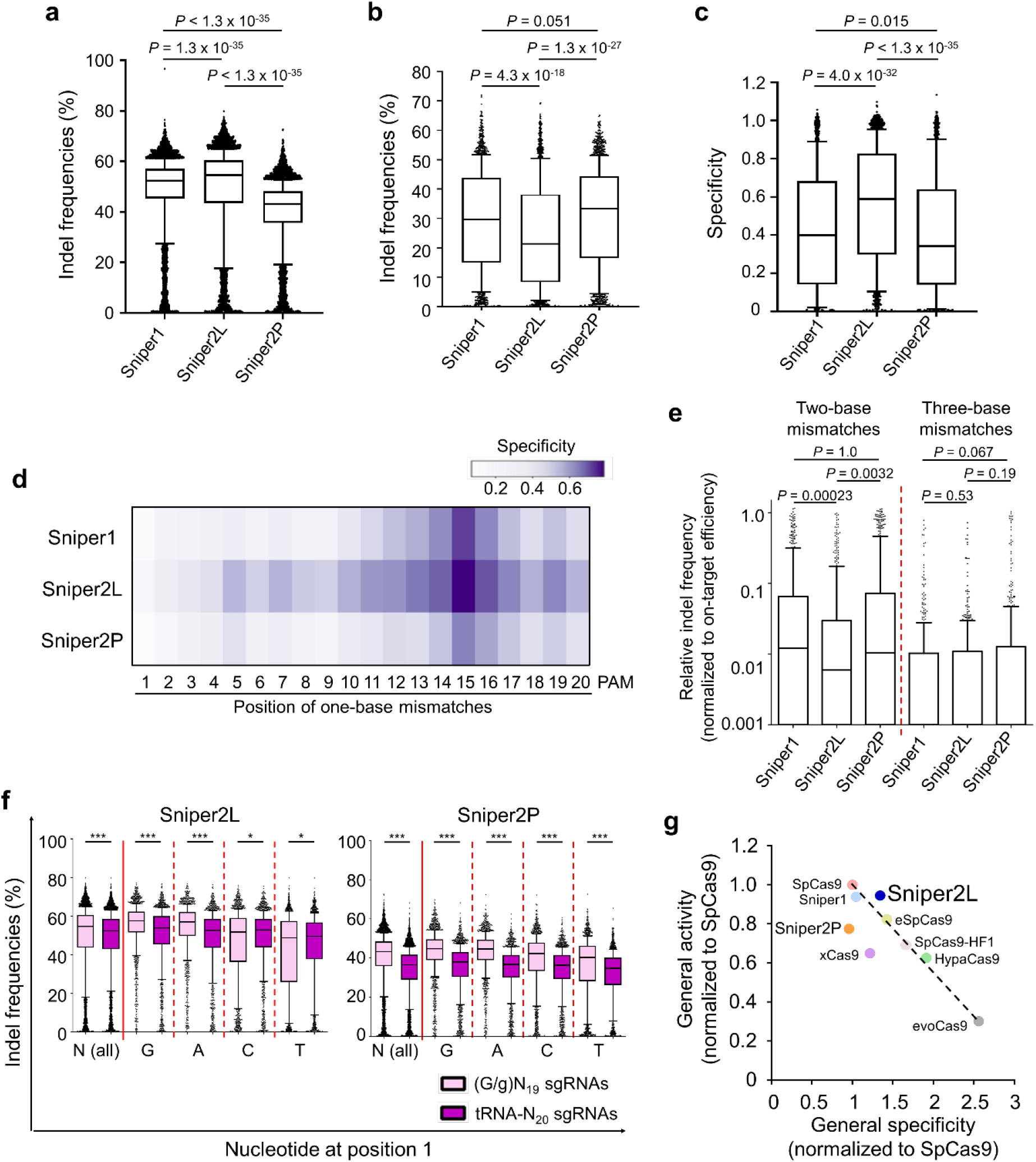
Comparison of Sniper-Cas9 variants. **a**, Indel frequencies at target sequences containing NGG PAMs. *n* = 7,702 target sequences. Kruskal-Wallis test. **b**, Indel frequencies at target sequences with single-base mismatches containing NGG PAMs. *n* = 1,732. Kruskal-Wallis test. **c**, General specificity of variants. Specificity was calculated as 1 – (indel frequencies at target sequences that harbor a single mismatch divided by those at perfectly matched target sequences). *n* = 1,734, 1,732, and 1,734 for Sniper1, Sniper2L, and Sniper2P, respectively. Mann-Whitney U test. **d**, Specificity of variants depending on the mismatch position (details in Supplementary Figure 10). **e**, Relative indel frequencies analyzed at target sequences with consecutive two- or three-base transversion mismatches. *n* = 554 and 531 for two- and three-base mismatches, respectively. The boxes represent the 25^th^, 50^th^, and 75^th^ percentiles; whiskers show the 10^th^ and 90^th^ percentiles. **f**, Activity assessments at target sequences with (G/g)N_19_ or tRNA-N_20_ sgRNAs. *n* = 6,321 (N), 1,666, 1,467, 1,626, and 1,562 (T) for Sniper2L, and 6,765 (N), 1,807, 1,587, 1,721, and 1,650 (T) for Sniper2P. *P* = 8.39 × 10^−20^ (N), 5.06 × 10^−26^ (G), 7.56 × 10^−34^ (A), 0.012 (C), and 0.04 (T) for Sniper2L, and 7.56 × 10^−34^ (all) for Sniper2P; Mann-Whitney U test. **g**, Relationship between the specificity and activity of SpCas9 and Cas9 variants. Sniper2L represents an outlier of the general trade-off. The specificity and activity of the high-fidelity variants were taken from our previous study^8^. The dashed line shows the general trade-off relationship.

Next, we compared the specificities of the Sniper-Cas9 variants with that of Sniper1 by comparing activities at mismatched target sequences using library A. Given that a comparison of activities at mismatched target sequences can be biased when the activities at matched target sequences are substantially different between the comparison groups, we used 30 sgRNAs that induced comparable Sniper-cas9 variant-directed indel frequencies either four or seven days after transduction (Supplementary Fig. 10). Each of the 30 sgRNAs was paired with 98 target sequences harboring one-, two-, or three-base mismatches (Methods). The activities of Sniper2L at the mismatched target sequences were significantly lower than those of Sniper1 and Sniper2P (Fig. 1b). If we define specificity as 1 – (indel frequencies at target sequences that harbor a single-mismatch divided by those at perfectly matched target sequences)^8^, the specificity of Sniper2L was significantly higher than those of Sniper2P and Sniper1 (Fig. 1c).

When we determined the specificity as a function of the mismatch position, we found that all three Sniper Cas9 variants showed higher specificity at the PAM-proximal region as compared to the PAM distal regions (Fig. 1d and Supplementary Fig. 11). Similar higher specificities at the PAM-proximal regions were also previously observed in other high-fidelity Cas9 variants^8^. Notably, Sniper2L was less likely to tolerate mismatches in both the PAM-distal and -proximal regions as compared to Sniper1 and Sniper2P; in those regions, local specificity reached high points at positions 5 and 15, respectively, which is compatible with the results of most previously reported high-fidelity variants^8^.

Furthermore, all Sniper variants tolerated single-base wobble mismatches more than single-base transversion mismatches (Supplementary Fig. 12), which is in line with the previous results of SpCas9 variants^8, 11^. The relative indel frequencies at mismatched target sequences containing two- or three-base transversion mismatches were dramatically reduced (Fig. 1e and Supplementary Fig. 13). Based on these results, we selected Sniper2L as our new version of Sniper-Cas9.

Since perfectly matched sgRNAs generated by the tRNA-associated processing system could increase the activity of some high-fidelity variants such as eSpCas9(1.1), SpCas9-HF1, and evoCas9, but not HypaCas9 and xCas9^8, 12, 13^, we compared the activities of the Sniper variants at identical targets using library C, based on tRNA-N_20_ sgRNAs, and library B, based on (G/g)N_19_ sgRNAs (hereafter, 20-nt guide sequences with a matched or mismatched 5′ guanosine are described as GN_19_ and gN_19_, respectively). Such (G/g)N_19_ sgRNAs are expressed from a U6 promoter with a G at the 5’ terminus, which is often mismatched with the corresponding nucleotide (position 1) at the target sequence. We observed that Sniper2L and Sniper2P displayed slightly higher general activities with (G/g)N_19_ sgRNAs than tRNA-N_20_ sgRNAs although tRNA-N_20_ showed slightly higher Sniper2L-induced activities than gN_19_ sgRNAs at target sequences starting with 5’-C or -T (Fig. 1f).

### Sniper2L represents an outlier to the trade-off relationship between general activity and specificity

We previously observed a trade-off between general activity and specificity of SpCas9 variants^8^; when a high-fidelity variant displayed high fidelity or specificity, it also exhibited relatively low general activity. To examine whether the Sniper2 variants followed this trend, we measured their activity and specificity using eight sgRNAs that were previously used in the analysis of the other high-fidelity variants. We observed that Sniper2L displayed both enhanced fidelity and higher on-target activities compared to Sniper1. To our knowledge, Sniper2L is the first and the only high-fidelity variant without sacrificing its general activity, being an outlier to the trade-off relationship between general activity and specificity (Fig. 1g).

### Computational models to predict activities of Sniper2L

Given that the activities of Sniper2L at matched and mismatched target sequences are dependent on the target sequence, accurate prediction of the Sniper2L activities would facilitate its utilization. Thus, we developed deep-learning based computational models that predict activities of Sniper2L and Sniper1 with (G/g)N_19_ and tRNA-N_20_ sgRNAs at matched target sequences (Fig. 2a and Supplementary Fig. 14a) and with (G/g)N_19_ sgRNAs at mismatched target sequences (Supplementary Fig. 14b). We randomly divided the data obtained from libraries A, B, and C into training and test data sets (Supplementary Tables 2-4). When we evaluated our models using the test data sets, we observed robust performance at both matched target sequences (Pearson correlation co-efficient r = 0.96, Spearman correlation co-efficient R = 0.94) and mismatched target sequences (r = 0.92, R = 0.90) (Fig. 2b,c). We collectively named these computational models DeepSniper and provided it as a web tool: http://deepcrispr.info/DeepSniper for wide use.

**Figure 2.**
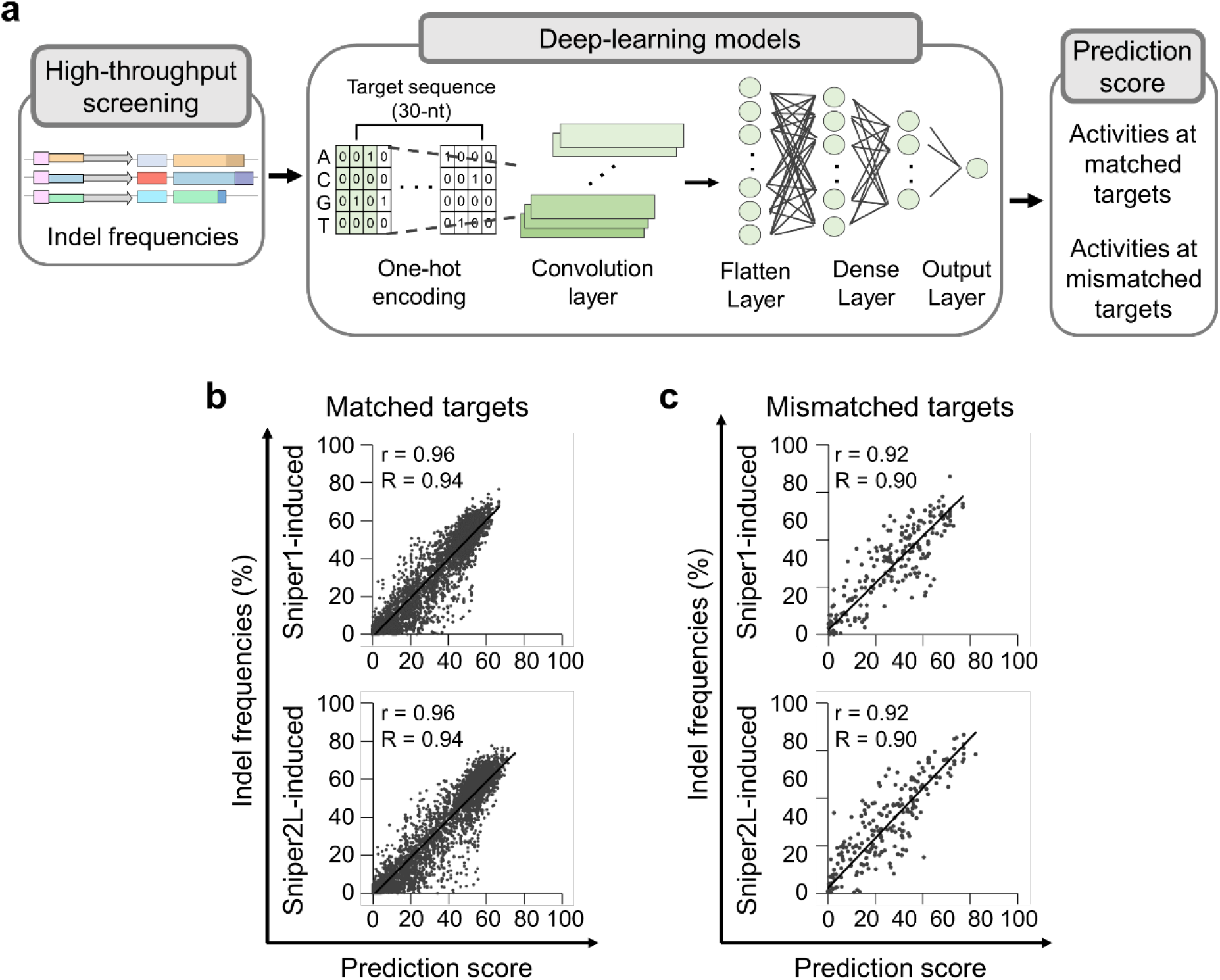
Development of deep-learning based prediction models, collectively named DeepSniper. **a**, A simplified schematic representation of DeepSniper development. **b**,**c**, Performance of DeepSniper in predicting the activities of Sniper1 and Sniper2L at matched (**b**) and mismatched (**c**) target sequences using target sequences that were not included in training data sets. The Pearson correlation coefficient (r) and the Spearman correlation (R) are presented. The number of target sequences *n* = 5,100 and 5,069 for Sniper1 and Sniper2L, respectively (**b**) and 295 for Sniper1 and Sniper2L (**c**).

### High-throughput evaluation of Cas9 variants delivered as RNPs

Cas9 and sgRNAs are frequently delivered preassembled RNP format during ex vivo genome editing therapy for human patients^14-16^. Given that delivery methods affect the on- and off-target activities of Cas9^17^, the results drawn from our analysis following lentiviral delivery might not be directly extrapolated to therapeutically relevant RNP-based approaches. Thus, we next attempted to measure the activities of high-fidelity variants, including Sniper2L, that had been delivered in RNP format into cells in a high-throughput manner (Supplementary Table 5). For this purpose, we utilized guide RNA swapping^18^ and our library of sgRNA and target sequence pairs. For accurate high-throughput evaluations, cells that do not express Cas9 protein should be removed and we used an antibiotic selection step to remove the cells that do not express Cas9^8, 19^. When the Cas9 delivery platform is changed from plasmid to RNP, this step is no longer available. To overcome this limitation, we delivered Cas9 protein together with an *HPRT*-targeting sgRNA. Because *HPRT* knockout provides resistance to 6-thioguanine (6-TG), the cells in which Cas9 delivery has occurred can be selected via 6-TG selection, similar to the antibiotic selection step (Fig. 3a).

**Figure 3.**
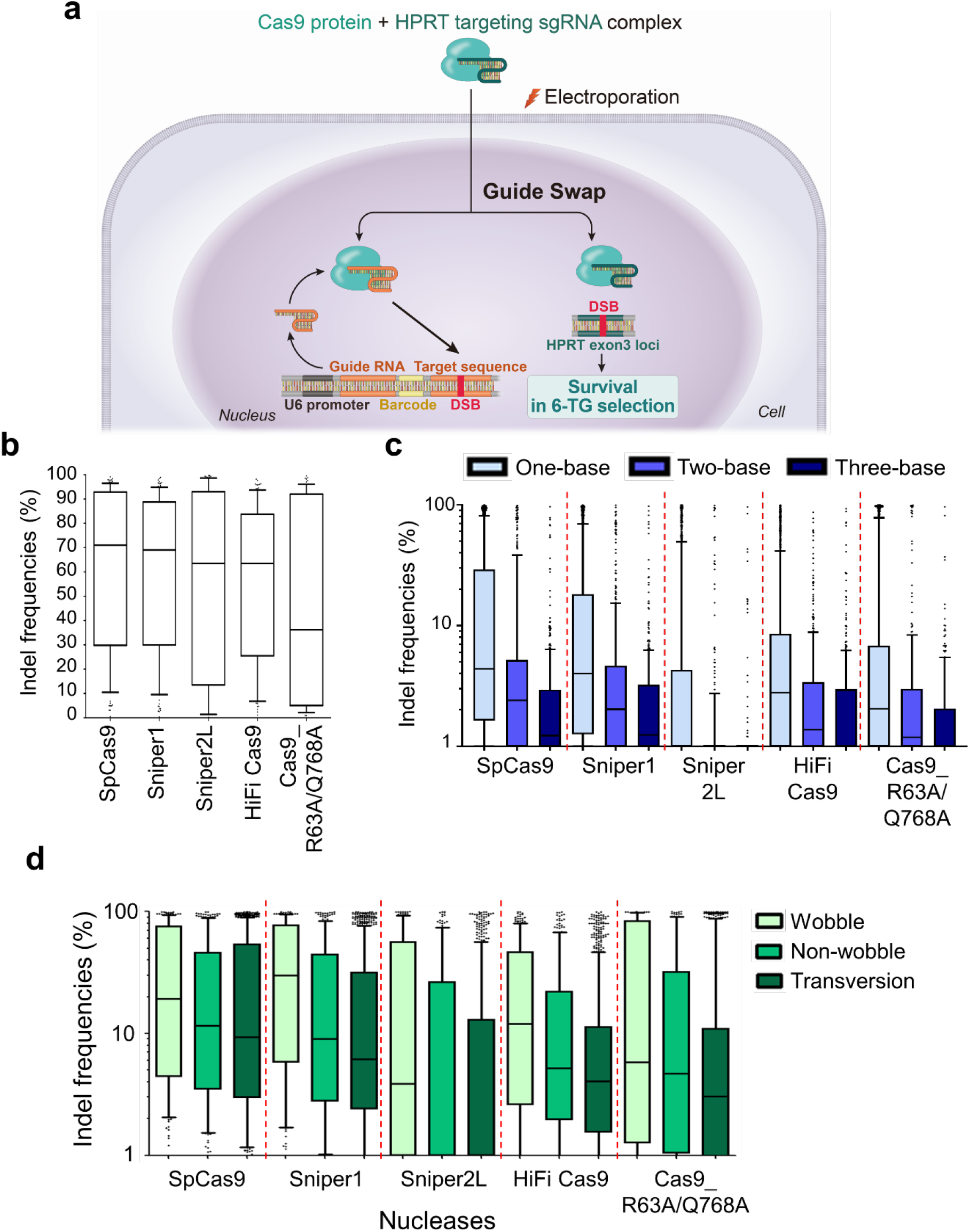
High-throughput evaluation of the activities of Cas9 variants when delivered as ribonucleoproteins. **a**, Schematic representation of experimental strategy using RNP delivery. **b**, Indel frequencies at perfectly matched target sequences with NGG PAMs. *n* = 105, 113, 81, 113, and 69 for SpCas9, Sniper1, Sniper2L, HiFi Cas9, and Cas9_R63A/Q768A, respectively. No statistically significant difference; Kruskal-Wallis test. **c**, Effects of the number of mismatches between the sgRNA and target sequence. *n* = 2,236 (one-base), 448 (two-base), and 414 (three-base) for SpCas9, 2,352, 446, and 296 for Sniper1, 1,641, 322, and 296 for Sniper2L, 2,248, 441, and 414 for HiFi Cas9, and 1,398, 278, and 245 for Cas9_R63A/Q768A. **d**, Activities of variants at target sequences with single-base mismatches as a function of the type of mismatch. *n* = 214 (wobble), 237 (non-wobble), and 923 (transversion) for SpCas9, 223, 246, and 982 for Sniper1, 169, 159, and 695 for Sniper2L, 223, 236, and 934 for HiFi Cas9, and 127, 144, and 604 for Cas9_R63A/Q768A. The boxes represent the 25^th^, 50^th^, and 75^th^ percentiles; whiskers show the 10^th^ and 90^th^ percentiles.

HEK293T cells were transduced with library A lentivirus at an MOI of 0.1. After puromycin selection to remove untransduced cells, we individually transfected SpCas9, Sniper1, Sniper2L, HiFi Cas9^6^, and Cas9_R63A/Q768A^4^, preassembled with the *HPRT*-targeting sgRNA, into the cell library. HiFi Cas9 and Cas9_R63A/Q768A were selected as the comparison groups because HiFi Cas9 showed low off-target effects when delivered as RNP format^6^ and because Cas9_R63A/Q768A is a very recently reported high-fidelity variants of SpCas9^4^. Then, we removed cells in which Cas9 were not delivered by 6-TG selection and isolated genomic DNA and analyzed it. As expected, 6-TG selection dramatically increased cells containing indels at *HPRT* (Supplementary Fig. 15), indicating that cells do not have Cas9 were removed. Some of these transfected Cas9 proteins pre-complexed with an *HPRT*-targeting sgRNA were expected to be complexed with a guide RNA expressed from the transduced library after guide swapping^18^ and then to target the corresponding target sequence in the library (Fig. 3a).

Given that such RNP-based high-throughput evaluation of Cas9 had not been conducted previously, we first determined the PAM sequences that were recognized by the high-fidelity variants to verify our strategy. Among target sequences containing all possible 4-nt PAM sequences (NNNN), variants displayed the highest indel frequencies at targets with NGG PAMs, which is in line with the results from Cas9 variant-expressing cell lines (Supplementary Fig. 16). However, we barely observed activities higher than 5% at target sequences containing non-canonical PAM sequences such as NGA or NAG. These results suggest that the shorter time of exposure to Cas9 (delivered in an RNP format) affected the efficiencies of the high-fidelity variants, such that they preferentially cleaved targets containing the most active PAM sequences.

We next assessed nuclease activities at 30 perfectly matched target sequences in library A and found that the activities of the high-fidelity variants were similar except that Cas9_R63A/Q768A showed a tendency of relatively lower activities, which is in line with the previous report^4^, although this difference was not statistically significant (Fig. 3b).

We also measured indel frequencies at mismatched target sequences and found that Sniper2L was highly inactive at the mismatched targets as compared to the other variants (Fig. 3c). Wobble single-base mismatches were more tolerated as compared to transversion mismatches for all variants (Fig. 3d). When we evaluated indel frequencies as a function of the mismatch position, Sniper2L hardly induced cleavage at target sequences with single-base or two- or three-base mismatches in PAM-proximal or -distal regions, a finding that is consistent with our results using lentivirus (Supplementary Fig. 17-19). Taken together, our results indicate that Sniper2L exhibits high on-target activities along with relatively low off-target activities compared to previously reported high-fidelity variants when delivered in either lentiviral or RNP format.

### Mechanistic evaluation of Cas9 variants using single-molecule DNA unwinding assay

We next determined the fidelity of Cas9 variants using a single-molecule approach^20^. Mechanistically, Cas9 first binds DNA via its recognition of PAM, directionally unwinds DNA protospacer from the PAM-proximal to the PAM-distal side while annealing the guide RNA to the target strand^21^ until ∼17 base pairs are unwound^22^ when Cas9 undergoes a major conformational change to activate its nucleases^23^, creating a double strand break. Mismatches hinder unwinding, giving Cas9 its sequence specificity^22^. High-fidelity Cas9 variants show higher sequence specificity in unwinding which we can quantify using the single-molecule FRET-based DNA unwinding assay^22, 24^.

We used a panel of DNA sequences that contain 0 to 4 consecutive PAM distal mismatches (Fig. 4a). The number of PAM distal mismatches, *n*_PD_, required for more than a two-fold decrease in fraction unwound, *f*_unwound_, was smaller for high fidelity variants (*N*_pd_ ≥ 3 for spCas9, *n*_PD_ ≥ 2 for Sniper1 and Sniper2P, and *n*_PD_ ≥ 1 for Sniper2L), making Sniper2L the most specific among them (Fig. 4b). Unwinding specificity, defined as 1 – (*f*_unwound_ for a target with a single-mismatch divided by *f*_unwound_ for a perfectly matched target), was also the highest for Sniper2L (Fig. 4c). We also tested a target sequence without or with a single mismatch in the 10^th^ position and found that Sniper2L gives a superior unwinding specificity of 0.83 compared to 0.33 of spCas9 (Fig. 4d).

**Figure 4.**
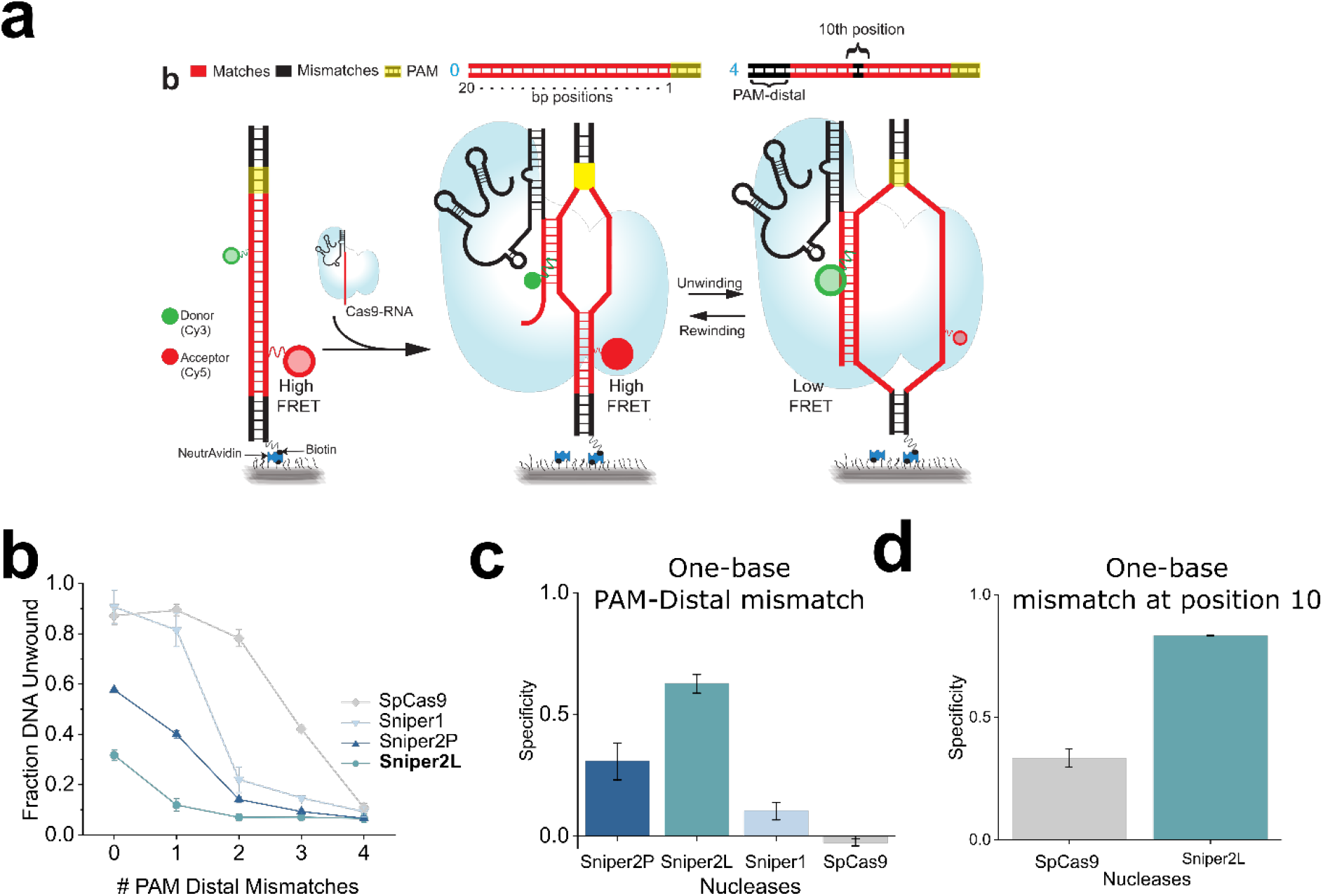
High unwinding specificity of Sniper2L revealed by single-molecule FRET. **a**, Schematic of a smFRET assay for investigation of Cas9-RNA induced DNA unwinding of surface immobilized DNA. DNA targets (top) are either a complete match to gRNAs (red) or contain mismatches (black) in the PAM distal (*n*_PD_) region or at position 10 relative to the gRNA. Unwinding increases the distance between donor and acceptor resulting in low FRET after > 10 bp of DNA are unwound (*E* 0.2 to *E* 0.6). **b**, *f*_unwound_ (equal to the relative fraction of molecules with *E of* 0.2 to *E* 0.6) vs. *n*_PD_ for different Cas9s. Error bars represent standard error (SE). n = 3 (or more) technical replicates. **c**, Unwinding specificity for different Cas9s calculated using a single PAM distal mismatch. Error bars represent SE. n = 3 (or more) technical replicates. **d**, Unwinding specificity for different Cas9s calculated using a mismatch at position 10. Error bars represent SE. n = 3 (or more) technical replicates.

## Discussion

In this study, we performed a directed evolution screen to generate Sniper2L, which was obtained through the addition of a further point mutation to Sniper-Cas9, a previously generated high-fidelity variant. Sniper2L was then characterized using two high-throughput evaluation methods, one involving lentiviral delivery and the other involving RNP delivery. Sniper2L showed higher specificity and higher general activity than Sniper-Cas9 and revealed higher specificity and comparable general activity as compared with SpCas9. Notably, this improvement shows that Sniper2L is an outlier to the previously found trade-off between general activity and specificity.

In addition, we developed a method for evaluating the activities of a large number of sgRNAs when Cas9 and sgRNA RNP complexes were delivered via electroporation. This new high-throughput methods are relevant to RNP delivery of Cas9 protein and sgRNAs, which is frequently used for ex vivo genome editing therapy for human patients. For successful clinical application of CRISPR technology, the selection of sgRNA with high activity and low off-target effects are crucial. For this purpose, researchers often have a large number of candidate sgRNAs and evaluating all of them often require a large amount of time and cost. In this case, our high-throughput evaluation method based on Cas9-sgRNA RNP complex delivery would facilitate the screening of sgRNAs.

While single-molecule unwinding analysis showed that Sniper2L has a superior discrimination against mismatched targets, its unwinding activity for a fully matched sequence was significantly lower compared to SpCas9 and Sniper1 for both of the DNA targets tested, suggesting that, for Sniper2L, the single-molecule unwinding readout does not accurately capture on target gene editing activities. We observed DNA molecules that are stably unwound or stably rewound at single molecule measurement time scale of ∼ minute. Single Sniper2L RNPs may come on and off the on-target site multiple times at gene editing scale of hours, yielding high gene editing activities.

In summary, by rounds of screening following random mutagenesis, we identified Sniper2L a new high-fidelity variant that exhibits an editing efficiency almost comparable to that of SpCas9, representing an outlier to the trade-off between general activity and specificity. We expect that Sniper2L will be very useful for genome editing when high-efficiency and low levels of -off-target effects are required.

## Acknowledgments

We would like to thank Younghye Kim and Seonmi Park for assisting with the experiments and Myungjae Song for providing technical advice. This work was supported in part by ‘Alchemist Project’ funded by the Ministry of Trade, Industry and Energy (grant no. 20012443 (to J.K.L.)), the National Research Foundation of Korea (NRF) grant funded by the Korea government (MSIT) (No. 2017M3A9B4062403 (to H.H.K.), 2018R1A5A2025079 (to H.H.K.), and 2022R1C1C2004229 (to N.K.)), the Yonsei Signature Research Cluster Program of 2021-22-0014 (to H.H.K.), the Brain Korea 21 FOUR project for Medical Science (Yonsei University College of Medicine), and the Korean Health Technology R&D Project, Ministry of Health and Welfare, Republic of Korea (grant HI21C1314 (to H.H.K)), and the US National Institutes of Health (grant R35 GM122569). TH is an investigator with the Howard Hughes Medical Institute. This article is subject to HHMI’s Open Access to Publications policy. HHMI lab heads have previously granted a nonexclusive CC BY 4.0 license to the public and a sublicensable license to HHMI in their research articles. Pursuant to those licenses, the author-accepted manuscript of this article can be made freely available under a CC BY 4.0 license immediately upon publication.

## Author contributions

YK, NK, JL, and KC performed the experiments. YK, NK, HHK, TH, and JKL designed the study and wrote the manuscript. SC and SM developed computational models. Y-HK and J-SK provided valuable advice during the project. IC, JC and VH performed single molecule measurements and analysis under supervision by TH.

## Competing financial interests

JL and JKL have submitted a patent disclosure on this work.

## Online Methods

### Plasmid construction

Each type of plasmid used in the Sniper-screen contains replication origins and resistance markers that are compatible with each other. The p11a plasmid, which contains the *ccdB* gene, was double-digested with SphI and XhoI enzymes (Enzynomics) and ligated to oligos (Cosmogenetech) containing the EMX1(1.6) target sequence (gcgccacTggttgatgtgat) with T4 DNA ligase (Enzynomics). The pSC101 (sgRNA-expressing vector) and the Sniper-Cas9 library plasmid have been described previously. The EMX(1.6) sgRNA sequence with a mismatch (gcgccacTggttgatgtgat; the mismatched nucleotide at position 13 is capitalized) was cloned into the pSC101 vector after BsaI digestion.

For generating plasmids that express Cas9 variants, the lentiCas9-Blast plasmid (Addgene, #52962) was digested with XbaI and BamHI-HF restriction enzymes (NEB, Ipswich, MA) and treated with 1 μl of calf intestinal alkaline phosphatase (NEB) for 30 min at 37 °C. The digested vector was gel purified using a MEGAquick-spin Total Fragment DNA Purification Kit (iNtRON Biotechnology, Seongnam, Republic of Korea) according to the manufacturer’s protocol. Mutation sites were introduced into variants by amplifying the lentiCas9-Blast plasmid using primers containing the mutation (Supplementary Table 6) with Phusion High-fidelity DNA Polymerase (NEB). The mutation sites were chosen according to suggestions from GenScript for inducing high variant expression levels^25, 26^.

The amplicons were gel-purified (iNtRON Biotechnology) and assembled with digested lentiCas9-Blast plasmids using NEBuilder HiFi DNA Assembly Master Mix (NEB) for 1 h at 50 °C.

### Sniper-Cas9 mutant library construction

Sniper-Cas9 mutant libraries were constructed using three independent protocols for mutagenesis, from XL1-red competent cells (Agilent), Genemorph II (Agilent), and Diversify PCR random mutagenesis (Clontech) kits. All reaction conditions have been described previously*. The assembled libraries were transformed into Endura™ electrocompetent cells (Lucigen) and incubated on LB plates containing chloramphenicol (12.5 μg/mL) at 37 °C overnight. A total of 3 × 10^6^ colonies were obtained for each library, resulting in an overall library complexity of 10^7^. Pooled library plasmids were purified using a midi prep kit (NucleoBond Xtra Midi EF, Macherey-Nagel).

### Positive and negative screening for directed evolution of Sniper-Cas9

BW25141-EMX1(1.6) was co-transformed with p11a (*ccdB* + target sequence) and pSC101 (sgRNA expression) plasmids (from which sgRNA expression can be induced by the addition of anhydrotetracycline (ATC)). The transformed BW25141-EMX1 cells were plated on LB plates containing ampicillin (50 μg/mL) and kanamycin (25 μg/mL), and then incubated overnight at 32 °C. Electrocompetent cells were produced from transformants cultured in liquid S.O.B. medium containing 0.1% glucose, ampicillin, and kanamycin until the OD600 reached 0.4. Each Sniper library underwent four round screening. 100ng of each library plasmid was transformed into 50 μL of electrocompetent BW25141-EMX1(1.6) cells using a Gene Pulser (Gene Pulser II, Bio-Rad) following the manufacturer’s instructions. In 1st screening, the transformed cell was incubated without ATC plated on chloramphenicol/kanamycin LB plates (nonselective conditions) and chloramphenicol/kanamycin/arabinose (1.5 mg/mL, Sigma-Aldrich) LB plates (selective conditions) without ATC followed by the overnight culture at 32 °C. In the 2nd to 4th screening, the transformed cell was incubated with 10 ng/ml ATC during recovery plated on nonselective and selective condition LB plates in the absence of ATC. Sniper screening conditions have been described previously*. After four rounds of screening, 50 colonies were obtained from selective condition plates, which were incubated in chloramphenicol containing LB medium at 42 °C. Each plasmid was Sanger-sequenced.

### Site saturation mutagenesis in Sniper-Cas9

For site saturation mutagenesis of the 1007^th^ position in Sniper-Cas9, the pBLC-Sniper-Cas9 plasmid was amplified using primers containing NNK at the appropriate position (Forward primer: agtaccccaagctggagagcnnkttcgtgtacggcgactacaagg; Reverse primer: tcttgatcagggcggtgcc). The PCR product was digested with DpnI (Enzynomics), treated with T4 polynucleotide kinase (Enzynomics), and ligated with T4 ligase (Enzynomics). The resulting product was transformed in DH5alpha cells. After Sanger-sequencing of plasmids from 100 randomly-selected colonies, variants containing 20 different amino acids at the 1007^th^ position were identified.

### Oligonucleotide libraries

Three oligonucleotide pools, libraries A, B, and C, were described in our previous study^8^. Library A was utilized for evaluating PAM sequences and activities at mismatched target sequences. Using library B, indel frequencies induced by variants were measured at a large number of target sequences with (G/g)N_19_ sgRNAs. Library C contained target sequences that were identical with those in library B but used a different sgRNA expression system that resulted in perfectly matched tRNA-N_20_ sgRNAs. All three oligonucleotide libraries were used for examining Sniper-Cas9 variants based on lentiviral delivery, whereas Library A was applied for comparing high-fidelity variants using the RNP delivery method.

### Cell culture and transfection

Human embryonic kidney 293T (HEK293T) cells were maintained in Dulbecco’s Modified Eagle Medium (DMEM) supplemented with 100 units ml^-1^ penicillin, 100 mg ml^-1^ streptomycin, and 10% fetal bovine serum. Cells were transfected using lipofectamine 2000 (Invitrogen) at a weight ratio of 1:1 (Sniper-Cas9 variant plasmid: sgRNA expression plasmid) in 48 wells. Genomic DNA was isolated with a DNeasy Blood & Tissue Kit (Qiagen) 72 h after transfection.

### Production of lentivirus

Lentivirus was produced using a method identical to that utilized in our previous study^8^. In brief, the day before transfection, HEK293T cells were seeded; the following day, the cells were treated with chloroquine diphosphate for up to 5 h, and transfected with lentiviral vector and packaging plasmids. The next day, the lentivirus-containing media was removed and fresh DMEM was added to the transfected HEK293T cells. The supernatant with viral particles was harvested 48 h after transfection; remaining library plasmids were degraded by treatment with Benzonase (Enzynomics, Daejeon, Republic of Korea) ^27, 28^.

### Generation of Sniper-Cas9 variant-expressing cell lines and transduction of lentiviral libraries

For measuring lentiviral titers, HEK293T cells were transduced with sequentially diluted aliquots of lentivirus-containing supernatant along with 10 μg ml^-1^ of polybrene and incubated overnight. The next day, both transduced and untransduced cells were treated with 20 μg ml^-1^ of blasticidin S (InvivoGen, San Diego, CA), and the number of surviving cells in the transduced population was counted when the untransduced cells were no longer viable^27^. Cell lines expressing Sniper-Cas9 variants were continuously maintained in the presence of 20 μg ml^-1^ of blasticidin S (InvivoGen).

Lentiviral libraries were transduced into Sniper-Cas9 variant-expressing cells using a protocol identical with that previously described^8^. In brief, 2.5 × 10^7^ Sniper-Cas9 variant-expressing cells were seeded in each 15-cm dish; two dishes (with a total of 5 × 10^7^ cells) were used for libraries A and C and four dishes (with a total of 1.0 × 10^8^ cells) were used for library B. Lentiviral plasmid libraries were transduced at an MOI of 0.4 along with 10 μg ml^-1^ of polybrene. After 4 days (libraries A, B, and C) and 7 days (library A) of transduction, cells were harvested.

### Western blotting

Levels of Sniper1.0, Sniper2L, and Sniper2P proteins were determined with Western blotting using purified anti-CRISPR Cas9 (diluted 1:1000, Biolegend, 844301) and anti-β-actin (diluted 1:1000, Santa Cruz Biotechnology, sc-47778) primary antibodies. Horseradish peroxidase-conjugated goat anti-mouse IgG antibody (diluted 1:5000, Santa Cruz Biotechnology, sc-516102) was used for signal detection.

### Deep sequencing and analysis

To examine the activities of the Sniper-Cas9 variants, samples were prepared and analyzed as previously described^8^. The following formula was used to remove background indel frequencies:

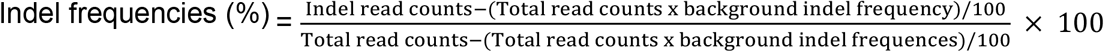

To minimize the errors generated by array synthesis, PCR amplification, or deep sequencing, we excluded target sequences with fewer than 100 total read counts or that exhibited background indel frequencies greater than 8% from the analysis.

### Deep learning models

Our data were randomly divided into training and test data sets, and 5-fold cross-validation was applied. For on-target prediction models, 32,109 and 31,810 target sequences were used for Sniper1 and Sniper2L, respectively (Supplementary Table 2-4). 2,656 and 2,654 target sequences were utilized for training the off-target prediction models for Sniper1 and Sniper2L, respectively (Supplementary Table 2). The numbers of target sequences that were used for evaluating the models are indicated in Figure 2b,c.

To develop on-target activity prediction models, the 30-nt target sequences were one-hot encoded to generate numerical inputs of the convolution layers, and zero-padding was utilized for retaining the number of target sequences. The features of the input sequences were extracted using the first convolution layer using 256 filters with 5-nt length for both Sniper1 and Sniper2L followed by average pooling layers, which were then flattened. Two fully connected layers with 1,500 nodes and one fully connected layer with 100 nodes were used for both Sniper1 and Sniper2L. To consider whether (G/g)N_19_ or tRNA-N_20_ sgRNA expression systems were adopted, they were indicated as a binary value. The features of a binary value were converted into a 100-dimensional vector and multiplied with the output of the third fully connected layer to integrate features of target sequence compositions and sgRNA expression systems. The final prediction scores were generated by performing a linear transformation of the output of the multiplication.

To develop off-target activity prediction models, the 20-nt sgRNA sequences and mismatched targets were one-hot encoded to make numerical inputs of the convolution layers, and zero-padding was used for sustaining the number of target sequences. The features of input sequences were extracted using the first convolution layer using 128 filters with 3-nt and 5-nt length for Sniper1 and Sniper2L, respectively, followed by average pooling layers, which were then flattened. As another input, the identities of mismatched nucleotides were given as numerical values, and those were concatenated with the output of the flatten layer. Three fully connected layers with 1,500 nodes and one fully connected layer with 100 nodes were utilized for both Sniper1 and Sniper2L, and the information about the sgRNA expression systems was not provided. The final prediction scores were generated by performing a linear transformation of the output of the multiplication.

Dropout layers with a rate of 0.3 were applied to avoid overfitting. The rectified linear unit (ReLu) was adopted for the convolution and dense layers. As the loss function, a mean absolute error was utilized, and an Adam optimizer with a learning rate of 10^−4^ was applied. TensorFlow was used for developing our models^29^.

### Delivery of Cas9 variants into a cell library using RNPs

Lentiviral plasmid library A was transduced into HEK293T cells at an MOI of 0.1 to generate a cell library. The cell library was continuously maintained in the presence of 2 μg ml^-1^ of puromycin (Invitrogen, Waltham, MA). The *HPRT*-targeting sgRNA templates were generated by annealing two complementary oligonucleotides, which were then incubated with T7 RNA polymerase in reaction buffer (40 mM Tris-HCl, 6 mM MgCl_2_, 10 mM DTT, 10 mM NaCl, 2 mM spermidine, 3.3mM NTPs, 1U/□ l RNase inhibitor, at pH 7.9) for 8 h at 37 °C. Transcribed sgRNAs were preincubated with DNase I to remove template DNA and purified using a PCR purification kit (Macrogen). A total of 3 × 10^7^ cells (6 × 10^6^ cells per dish x 5 dishes) were transfected with protein variants (WT-Cas9, Sniper1, Sniper2L, HiFi Cas9, and Cas9_R63A/Q768A; 40 μg) premixed with *in vitro* transcribed *HPRT*-targeting sgRNA (40 μg) and Alt-R Cas9 electroporation enhancer (4 μM, Integrated DNA Technologies) using a Neon transfection system (ThermoFisher) with the following settings: 1,150 V, 20 ms, and 2 pulses per 2 × 10^6^ cells using a 100 μL tip. On day 3 after transfection, a portion of the cell culture was harvested for analysis of indels at the *HRPT* site. Beginning on day 7 after transfection, cells were maintained with DMEM supplemented with 10% fetal bovine serum and 30 μM 6TG (Sigma). The cells were harvested 14 days after the 6TG selection began. Genomic DNA was isolated with a Blood & Cell Culture DNA Maxi Kit (Qiagen).

### Preparation of DNA targets for single molecule experiments

Integrated DNA Technologies (IDT) supplied all DNA oligonucleotides. For introducing Cy3 and Cy5 labels at the indicated locations, the oligonucleotides were synthesized with amine-containing modified thymine at the indicated location. A C6 linker (amino-dT) was used to label the DNA strands with Cy3 or Cy5 N-hydroxysuccinimido (NHS). For preparing the DNA, the non-target strand, target strand, and a 22 nt biotinylated adaptor strand were first mixed in 10 mM Tris-HCl, pH 8 and 50 mM NaCl. The mixture was transferred to a heat block preheated to 90 °C. After 2 minutes of heating, the mixture was cooled to room temperature over a few hrs. The sequences of the target and non-target strand (with same label positions) were changed to create DNA targets with mismatches. The full sequence of all DNA targets used in the smFRET assay is shown in Supplementary **Table 7**.

### Preparation of gRNA and Cas9-gRNA for single molecule experiments

crRNA and tracrRNA were synthesized by Integrated DNA technologies (IDT). All gRNAs were prepared by mixing crRNA (10uM) and tracrRNA (12uM) in 1:1.2 ratio in 10 mM Tris HCl (pH 8) and 50 mM NaCl. This mixture was then placed in a heating block pre-heated to 90 °C for 2 minutes and was allowed to cool to room temperature over a few hours for efficient hybridization between crRNA and tracrRNA. Cas9-RNA was prepared by mixing the gRNA (1uM) and Cas9 (2uM) at a ratio of 1:2 in Cas9-RNA activity buffer of 20 mM Tris HCl (pH 8), 100 mM KCl, 5 mM MgCl_2_, 5% v/v glycerol (final conc. 500nM). The full sequences of all the gRNA used in this study are available in Supplementary **Table 7**.

### Single-molecule fluorescence imaging and data analysis

Flow chamber surfaces coated with polyethylene glycol (PEG) was used for immobilization of DNA targets. These flow chambers were purchased from Johns Hopkins University Microscope Supplies Core. The neutrAvidin-biotin interaction was used for immobilizing the biotinylated DNA target molecules on the PEG-passivated flow chamber in the Cas9-RNA activity and imaging buffer without glucose oxidase and catalase (20 mM Tris-HCl, 100 mM KCl, 5 mM MgCl_2_, 5% (v/v) glycerol, 0.2 mg ml^-1^ BSA, 0.8% dextrose and saturated Trolox (>5 mM))(20). Cas9-RNA in the Cas9-RNA activity and imaging buffer with catalase and glucose oxidase (20 mM Tris-HCl, 100 mM KCl, 5 mM MgCl_2_, 5% (v/v) glycerol, 0.2 mg ml^-1^ BSA, 1 mg ml^-1^ glucose oxidase, 0.04 mg ml^-1^ catalase, 0.8% dextrose and saturated Trolox (>5 mM)) was added to the flow chamber at the concentrations much higher (E.g., 100 nM) than the dissociation constant of Cas9-RNA-DNA for Cas9-RNA targeting of DNA and Cas9-RNA induced DNA unwinding. All the imaging experiments were done at room temperature and time resolution was either 100 ms or 35 ms per frame. The total fluorescence from each of the immobilized DNA targets molecules was optically split into two separate donor and acceptor optical paths. The emissions belonging to these parts were projected onto two halves of a cryo-cooled (< -70 °C) EMCCD camera (Andor) which was stored as a video recording by the camera. The video recording containing fluorescent spots was then analyzed using custom scripts to extract background corrected donor fluorescence (I_D_), acceptor fluorescence (I_A_). FRET efficiency (*E*) of each detected spot was approximated as *E* = *I*_A_/(*I*_D_+*I*_A_). In the analysis of DNA unwinding experiments, the DNA molecules with the missing or inactive acceptor label were avoided by only including the fluorescent spots in the acceptor channel. The data acquisition software and analysis scripts can be downloaded from GitHub (https://github.com/Ha-SingleMoleculeLab). A detailed explanation of smFRET data acquisition and analysis has previously been described (Joo & Ha, *Cold Spring Harbor Protocols*, 2008).

### *E* histograms and analysis of Cas9-RNA induced DNA unwinding and rewinding

For every single molecule, the first five data points of its *E* time-traces were used as data points to construct *E* histograms. More than 2000 molecules contributed to each *E* histogram. The donor only peak (*E*=0), low FRET (0.2 <*E* <0.6 or 0.65 or 0.70) and high FRET (*E* >0.6 or 0.65 or 0.7) are three characteristic populations observed in these *E* histograms. Based on this low and high FRET populations, Cas9-RNA induced DNA unwinding was modeled as a two-state system, as shown below. The unwound fraction (*f*_unwound_) was calculated as a fraction of the low FRET population in the *E* histograms of DNA unwinding experiments.

## Statistical significance

Results from the Kruskal-Wallis test and Mann-Whitney U test calculated by SPSS Statistics (version 25, IBM) are shown.

## Reporting summary

Further information on the research design is available in the Nature Research Reporting Summary linked to this article.

## Data availability

We have submitted the deep sequencing data from this study to the NCBI Seqeuence Read Archive under accession number PRJNA817000. We have provided the data sets used in this study as Supplementary Tables 1-7.

## Code availability

We have made the source code for DeepSniper available on Github at https://github.com/NahyeKim/DeepSniper.

