## Supplementary Information for "Sniper2L, a high-fidelity Cas9 variant with high activity"

Contents:

Supplementary Figures 1 – 19

Supplementary Tables 1 – 7

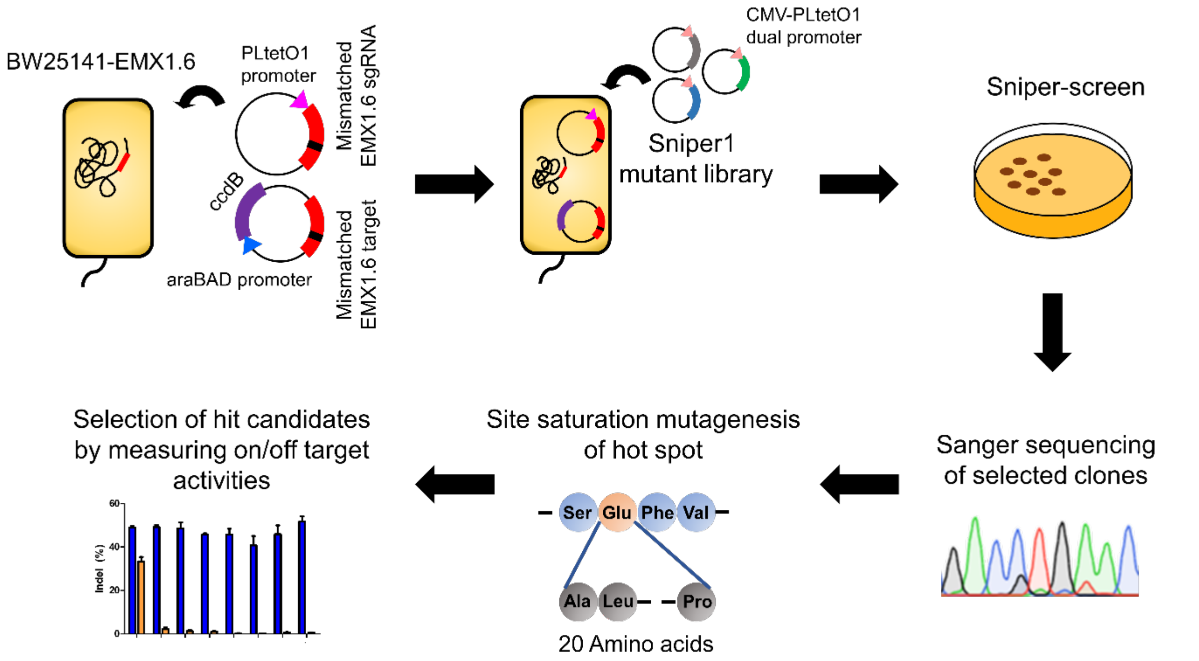


**Supplementary Figure 1.** Schematics of Sniper-screen and site saturation mutagenesis.

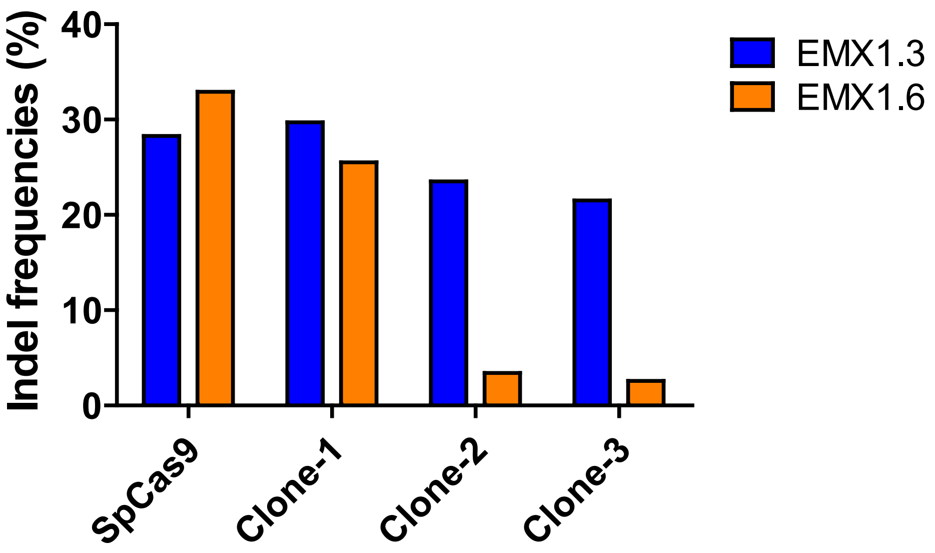


**Supplementary Figure 2.** Indel frequencies induced by Clone-1 (Sniper-Cas9, referred to in this manuscript as Sniper1.0), Clone-2, and Clone-3 with the EMX1.3 and EMX1.6 sgRNAs, which target two different sites in the human *EMX1* gene.

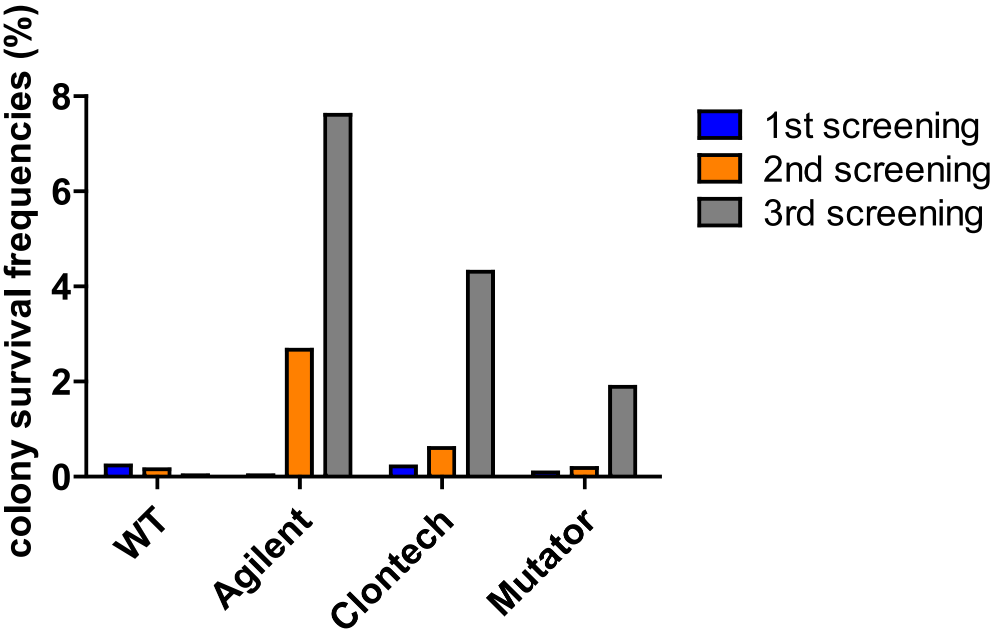


**Supplementary Figure 3.** Colony survival frequencies for cells transformed with three libraries encoding mutant versions of Sniper-Cas9, each generated by a different method of mutagenesis, after the first three rounds of the Sniper-screen. Plasmids derived from clones from the fourth screening were sequenced without colony counting to prevent contamination. Agilent, Genemorph II error-prone PCR kit from Agilent; Clontech, Diversify PCR random mutagenesis kit from Clontech; Mutator, XL-1 Red competent cells from Agilent.

| Amino acid, position of WT-Cas9 | Ag1 | Cl1 | Cl2 | Cl3 | Cl4 | Cl5 | Mu1 | Mu2 |
|---|---|---|---|---|---|---|---|---|
| **Lys, 4** | Lys | | | | | | | |
| **Lys, 112** | Asn | | | | | | | |
| **His, 137** | | His | | | | | | |
| **Ile, 350** | | Val | | | | | | |
| **Ile, 492** | Phe | | | | | | | |
| **Arg, 671** | His | | | | | | | |
| **Gln, 709** | | Gln | | | | | | |
| **Lys, 735** | Thr | | | | | | | |
| **Ala, 889** | | | | | | | | Val |
| **Glu, 1007** | | **Val** | | **Val** | **Gly** | **Gly** | | |
| **Tyr, 1021** | | Cys | | | | | | |
| **Lys, 1191** | | | | | | | Glu | |
| **Lys, 1192** | | | | | | Arg | | |
| **Ser, 1277** | | | Gly | | | | | |
| **Number of colonies for the clone/ Number of total sequenced colonies** | **11/12** | **14/39** | **8/39** | **3/39** | **3/39** | **2/39** | **4/48** | **3/48** |

**Supplementary Figure 4.** Sequencing results of selected hits obtained from the Sniper-screen performed with three classes of libraries encoding mutant versions of Sniper-Cas9. Ag1, Cl1~5, and Mu1~2 respectively indicate selected hits from libraries generated using the Genemorph II error-prone PCR kit, the Diversify PCR random mutagenesis kit, and XL-1 Red competent cells.

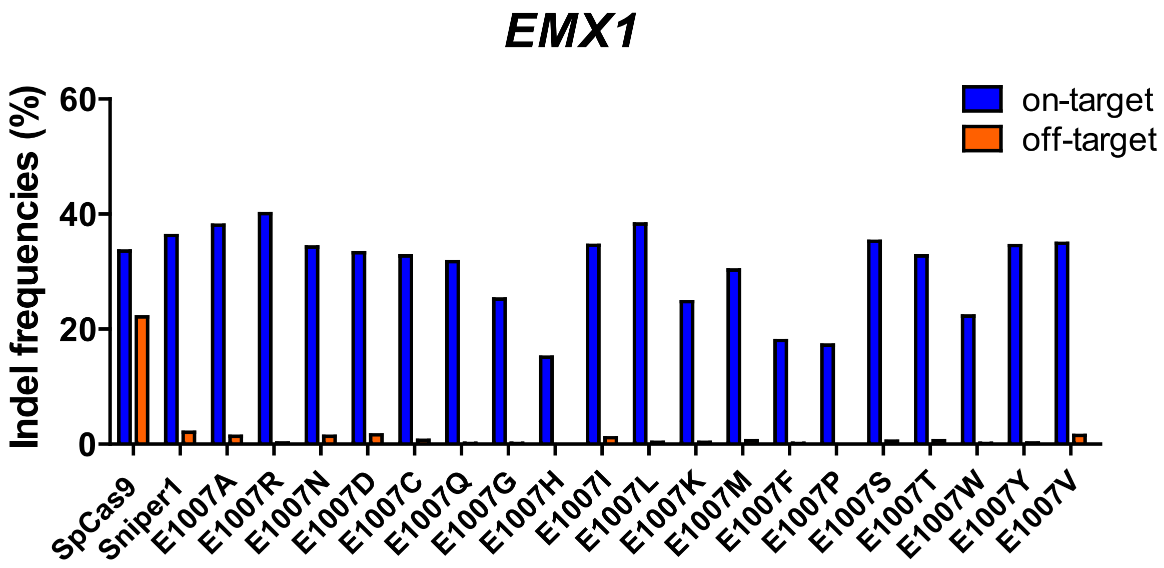


**Supplementary Figure 5.** Indel frequencies induced by SpCas9 and Sniper-Cas9 variants at matched and mismatched target sequences in HEK293T cells. Sniper-Cas9 variants were generated by site saturation mutagenesis at the 1007$^{th}$ amino acid (originally a Glu codon); the resulting amino acids at that position are shown on the x-axis. As the target sequence, a sequence (not EMX1.6 sgRNA-corresponding sequence) in the human *EMX1* gene was used.

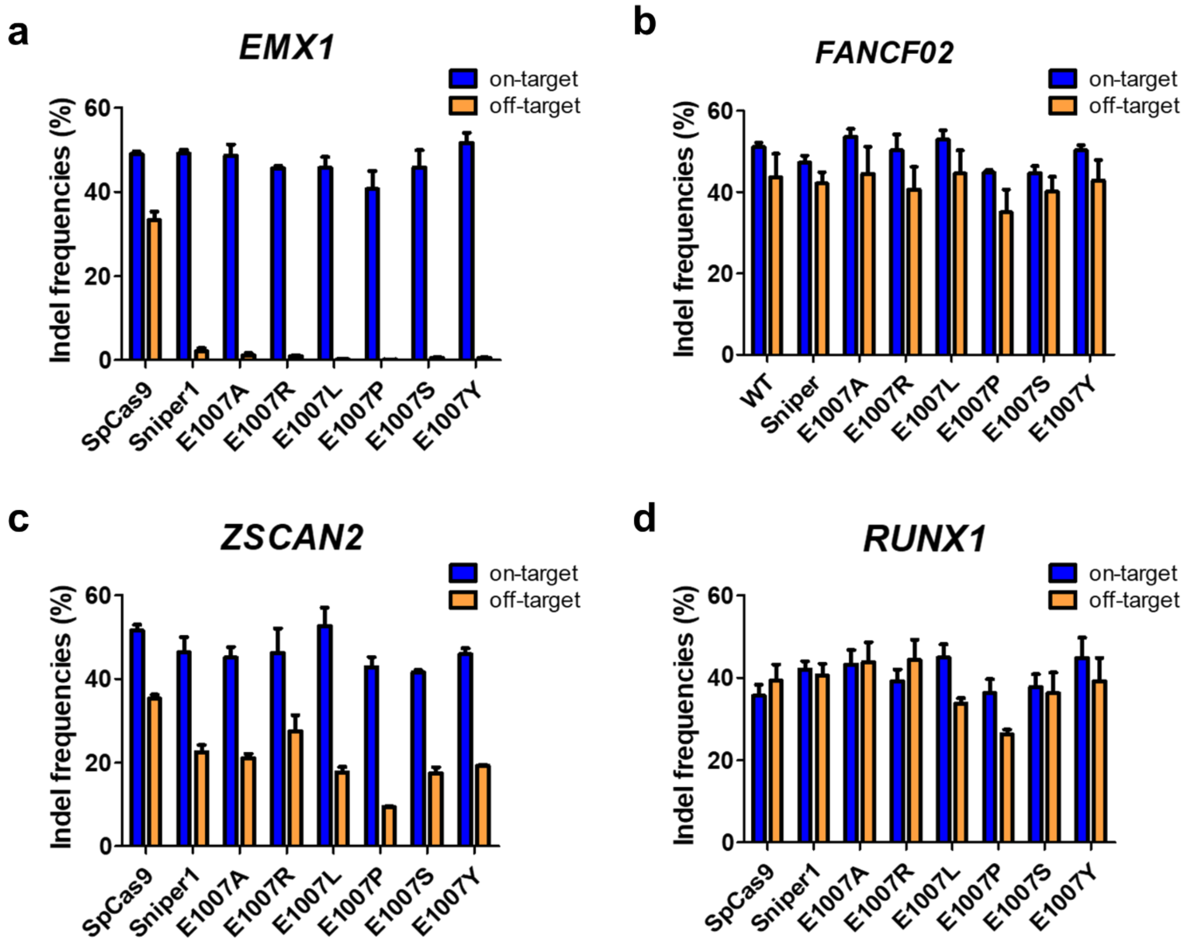


**Supplementary Figure 6.** Indel frequencies induced by SpCas9 and Sniper-Cas9 variants at matched and mismatched target sequences in HEK293T cells. Error bars indicate s.e.m. The number of independent transfection *n* = 3.

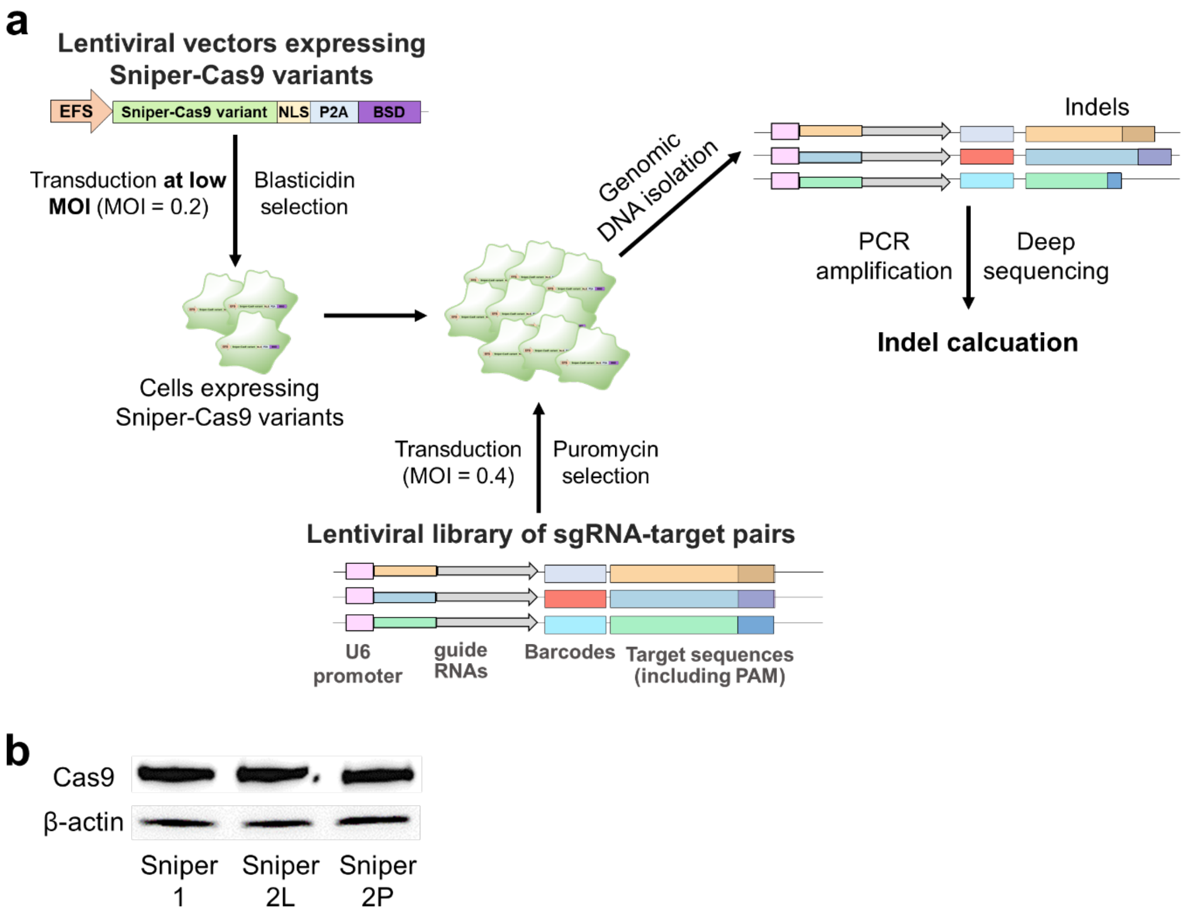


**Supplementary Figure 7.** Generation and examination of Sniper-Cas9 variant-expressing cell lines. **a**, Schematic representation of the generation of cell lines expressing Sniper-Cas9 variants and the subsequent evaluation of Sniper-Cas9 variants at a large number of target sequences. **b**, Western blot analysis to determine the level of expressed Cas9 proteins in the Sniper-Cas9 variant-expressing cell lines.

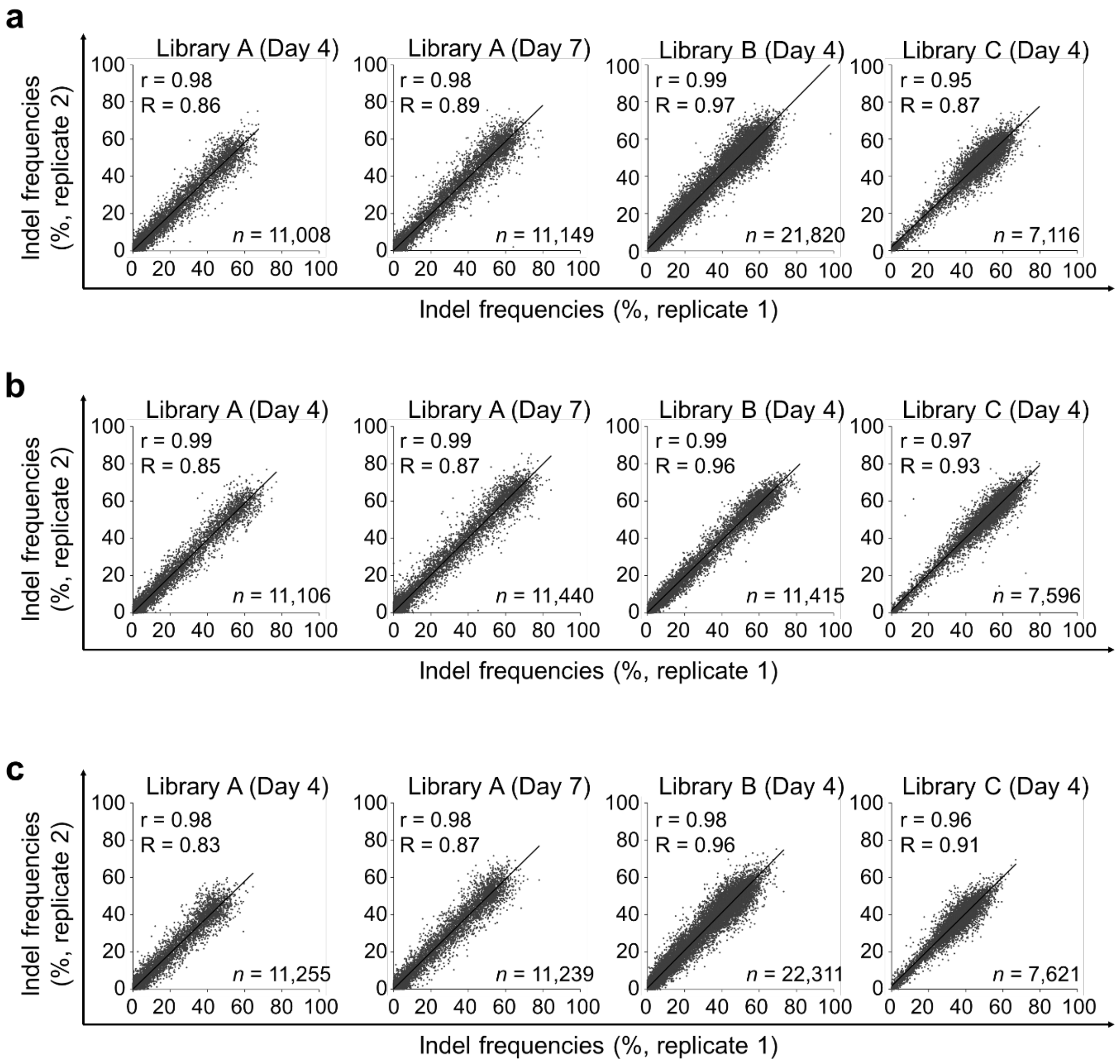


**Supplementary Figure 8.** Correlations between indel frequencies induced in two technical replicates by Sniper-Cas9 (**a**), Sniper2L (**b**), and Sniper2P (**c**) in the high-throughput analysis. The Pearson correlation coefficient (r) and the Spearman correlation (R) are shown. In each graph, the number of target sequences (*n*) that we used for analysis and the analyzed time (day) after the transduction of pair-wise libraries A, B, or C are shown.

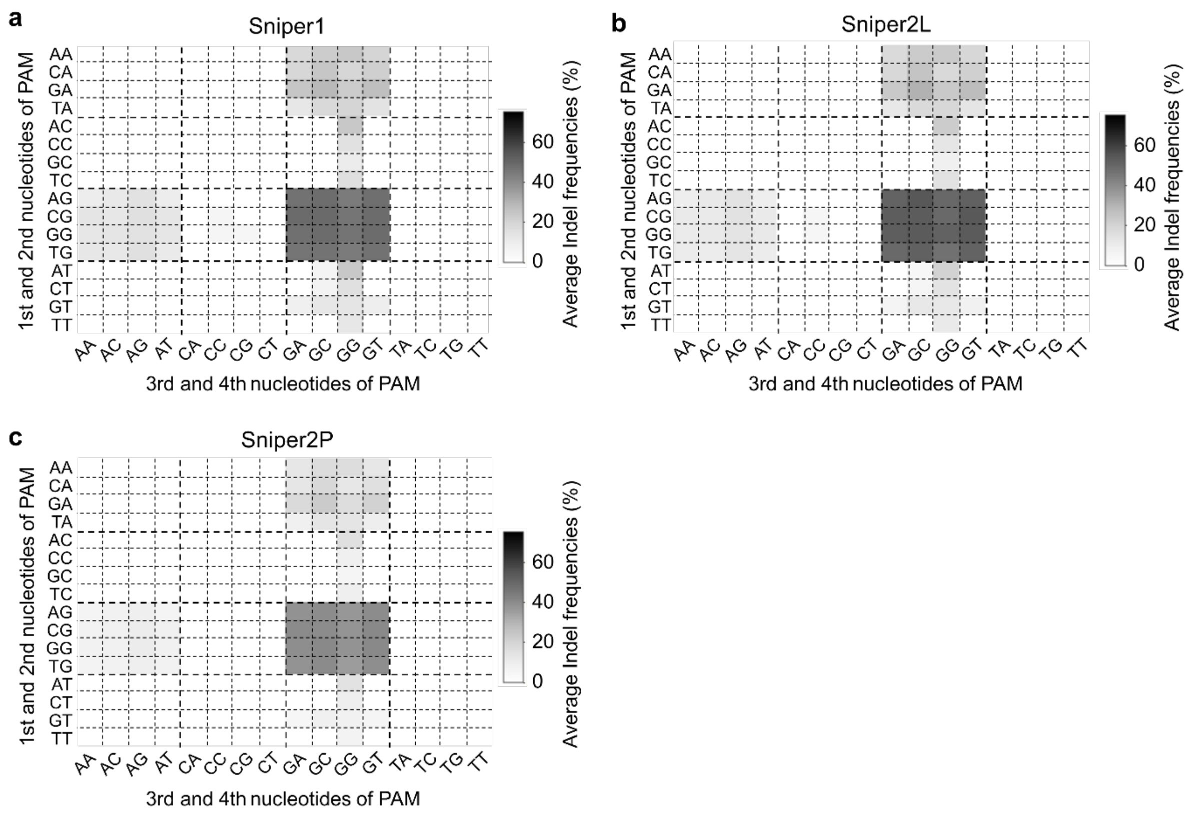


**Supplementary Figure 9.** PAM sequences recognized by Sniper1 (**a**), Sniper2L (**b**), and Sniper2P (**c**). Average indel frequencies four days after the transduction of library A into Sniper-Cas9 variant-expressing cells are shown; Average indel frequencies lower than 5% are indicated as white boxes in the grids. The number of target sequence *n* = 24 - 30 per PAM sequence.

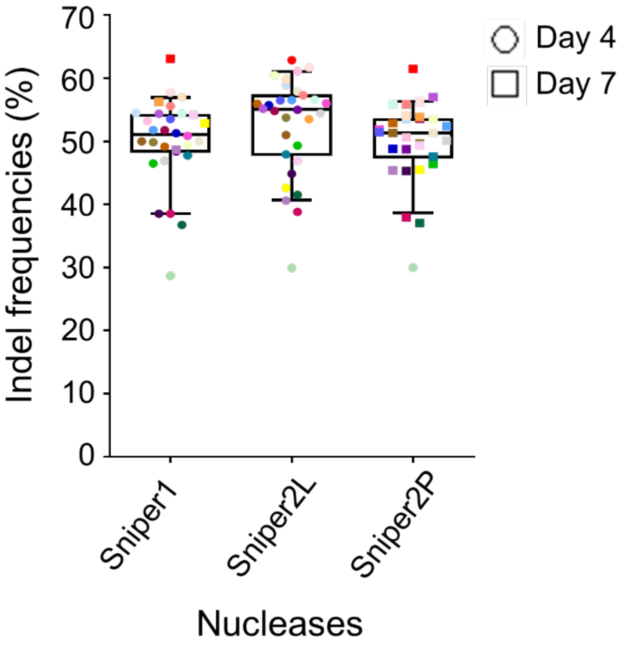


**Supplementary Figure 10.** Indel frequencies induced by Sniper1, Sniper2L, and Sniper2P measured using 30 perfectly matched guide RNA and target sequence pairs in library A on day 4 or 7 after transduction. The boxes represent the 25$^{th}$, 50$^{th}$, and 75$^{th}$ percentiles; whiskers show the 10$^{th}$ and 90$^{th}$ percentiles. The number of target sequence $n$ = 30. There is no statistically significant difference between results from the three variants; Kruskal-Wallis test.

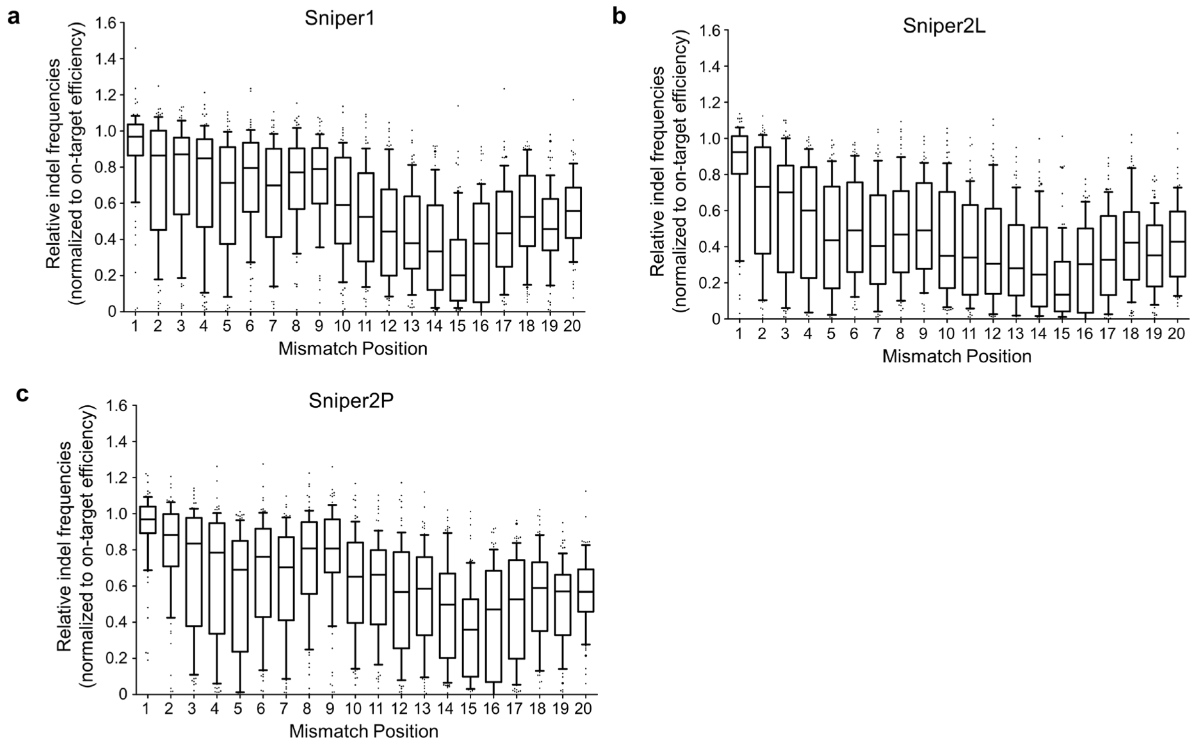


**Supplementary Figure 11.** Tolerance of Sniper1, Sniper2L, and Sniper2P for single-base mismatches varies depending on the mismatch position. The boxes represent the 25th, 50th, and 75th percentiles; whiskers show the 10th and 90th percentiles. The number of analyzed target sequence $n$ = 86 (position 1), 89 (position 2), 86, 87, 87, 87, 90, 83, 90, 90, 87, 86, 87, 89, 86, 86, 87, 85, 87, and 79 (position 20) for (**a**) and (**c**) and 86 (position 1), 89, 86, 86, 87, 87, 90, 83, 90, 90, 87, 86, 87, 89, 86, 86, 87, 84, 87, and 79 (position 20) for (**b**).

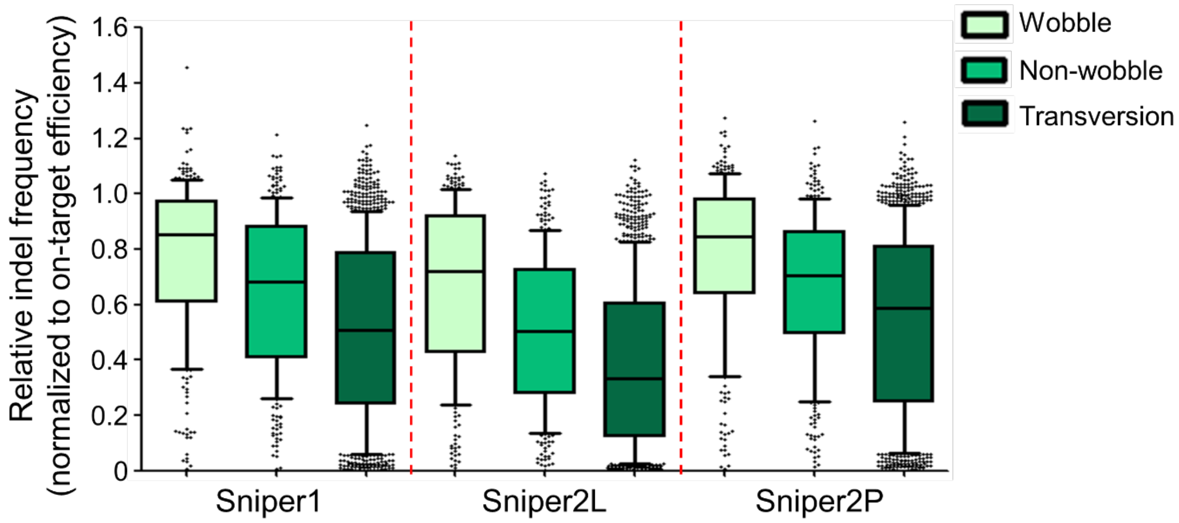


**Supplementary Figure 12.** Relative indel frequencies induced by Sniper1, Sniper2L, and Sniper2P at mismatched targets vary depending on the types of single-base mismatches. The boxes represent the 25th, 50th, and 75th percentiles; whiskers show the 10th and 90th percentiles. *n* = 275 (wobble), 304 (non-wobble), and 1,155 (transversion) for Sniper1 and Sniper2P and 275, 304, and 1,153 for Sniper2L.

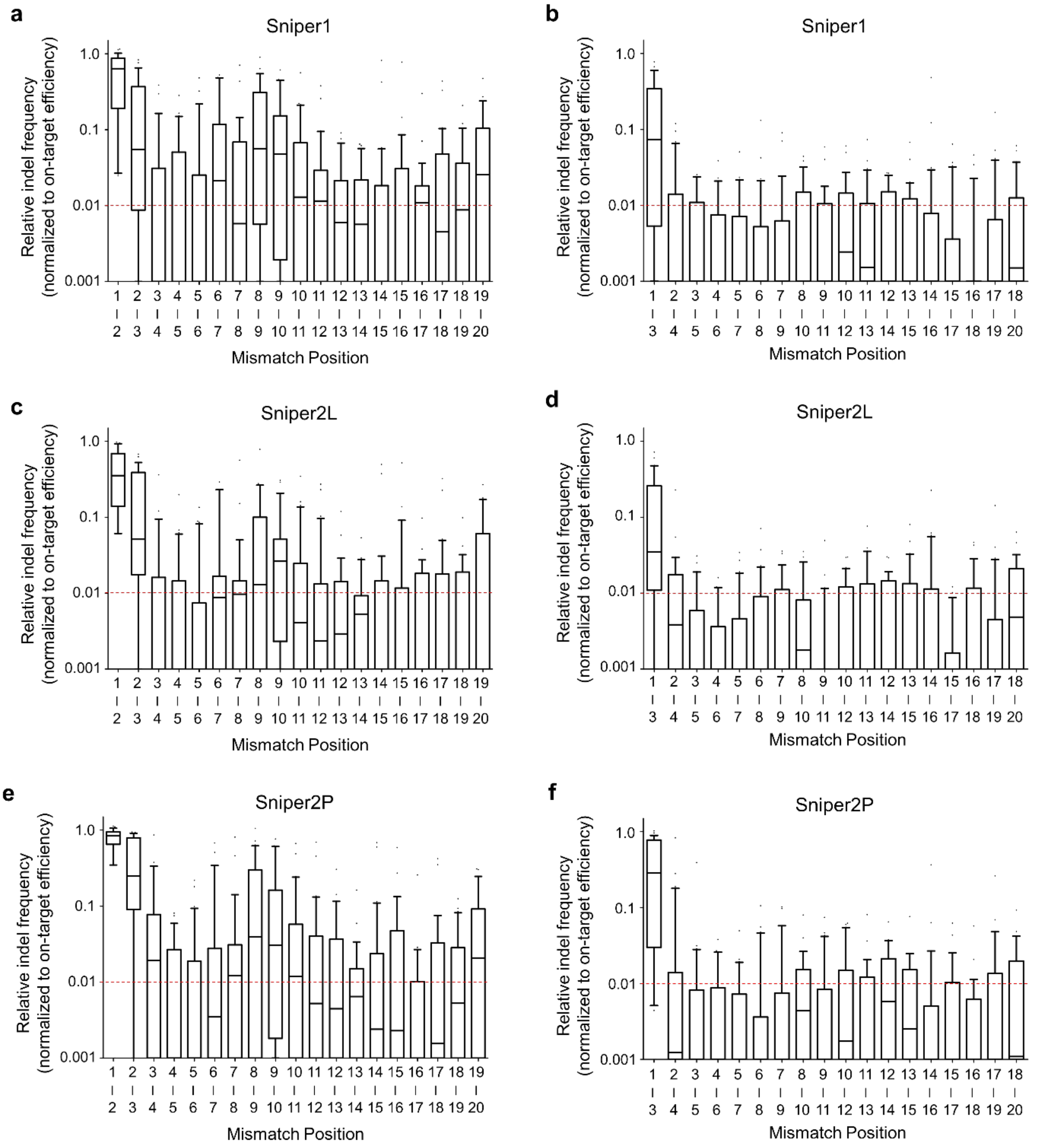


**Supplementary Figure 13.** Relative indel frequencies induced by Sniper1 (**a, b**), Sniper2L (**c**, **d**), and Sniper2P (**e**, **f**) at target sequences containing consecutive two- (**a**, **c**, **e**) or three- (**b**, **d**, **f**) base mismatches vary depending on the mismatch position. The boxes represent the 25$^{th}$, 50$^{th}$, and 75$^{th}$ percentiles; whiskers show the 10$^{th}$ and 90$^{th}$ percentiles. *n* = 30 (position 1-2), 29, 27, 30, 30, 29, 29, 30, 29, 30, 30, 29, 28, 30, 29, 28, 30, 30, and 27 (position 19-20) (**a**, **c**, **e**) and 30 (position 1-3), 30, 30, 29, 30, 30, 28, 28, 30, 30, 30, 28, 30, 30, 30, 29, 30, and 29 (position 18-20) (**b**, **d**, **f**).

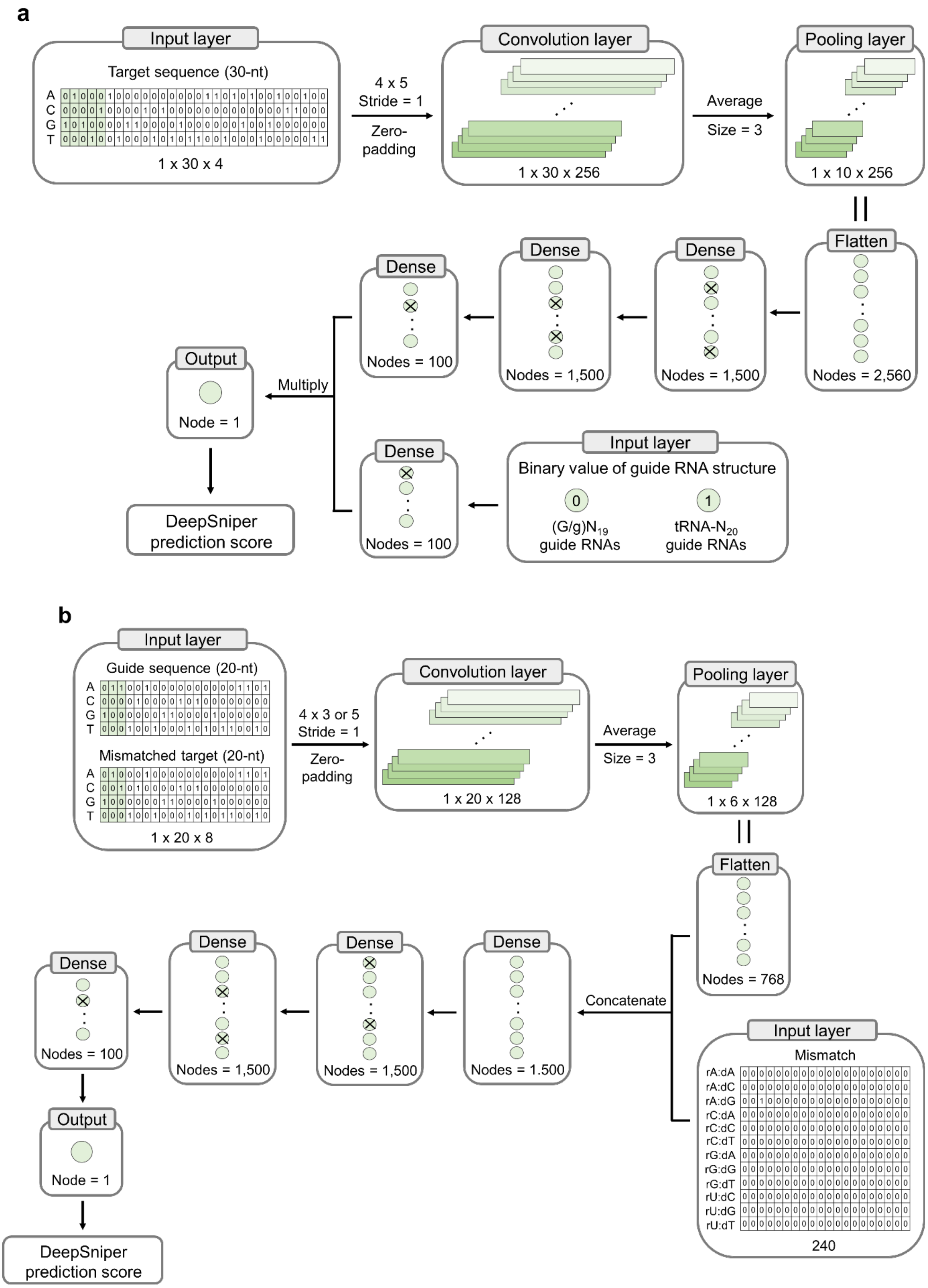


**Supplementary Figure 14.** Overview of DeepSniper, which predicts the on-target (**a**) and off-target (**b**) activities of Sniper-Cas9 variants.

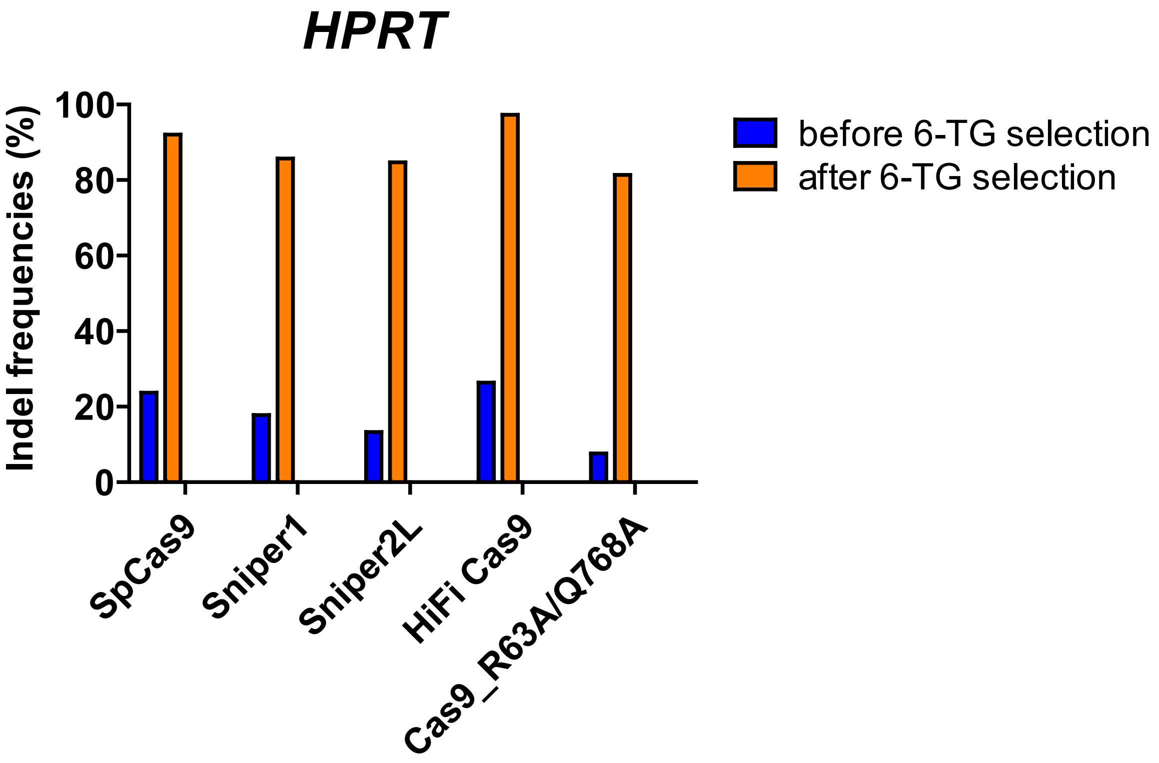


**Supplementary Figure 15.** Indel frequencies induced by Cas9 variants in the human *HPRT* gene in library screening using RNP delivery before and after 6-TG selection.

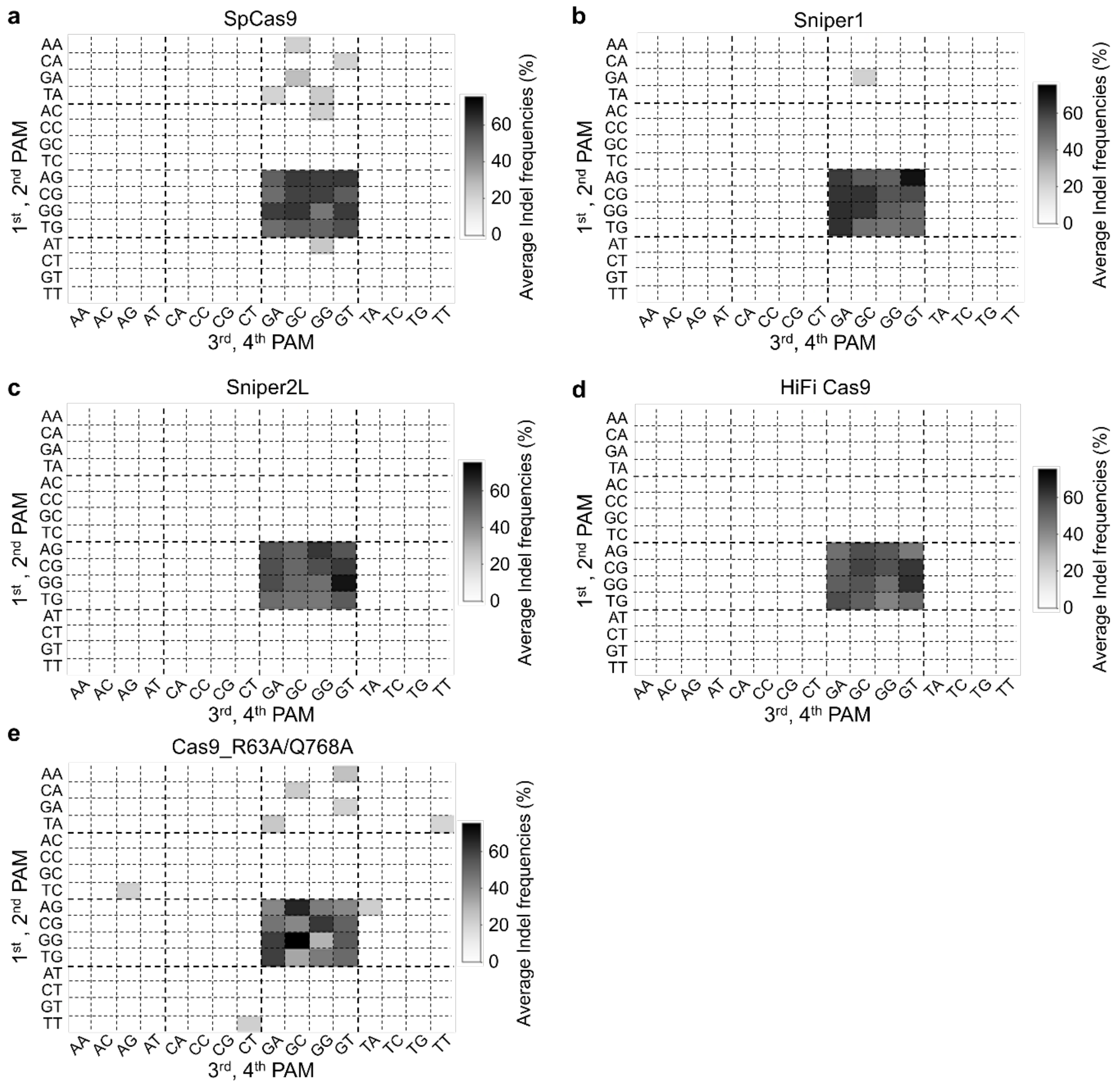


**Supplementary Figure 16.** Average indel frequencies associated with all possible 4-nt PAM sequences. We excluded PAM sequences associated with indel frequencies less than 5%; such PAMs are indicated as white boxes in the grid. The number of target sequence per each 4-nt PAM *n* = 14 – 28 (**a**), 15 – 29 (**b**), 12 – 28 (**c**), 13 – 29 (**d**), and 7 – 26 (**e**).

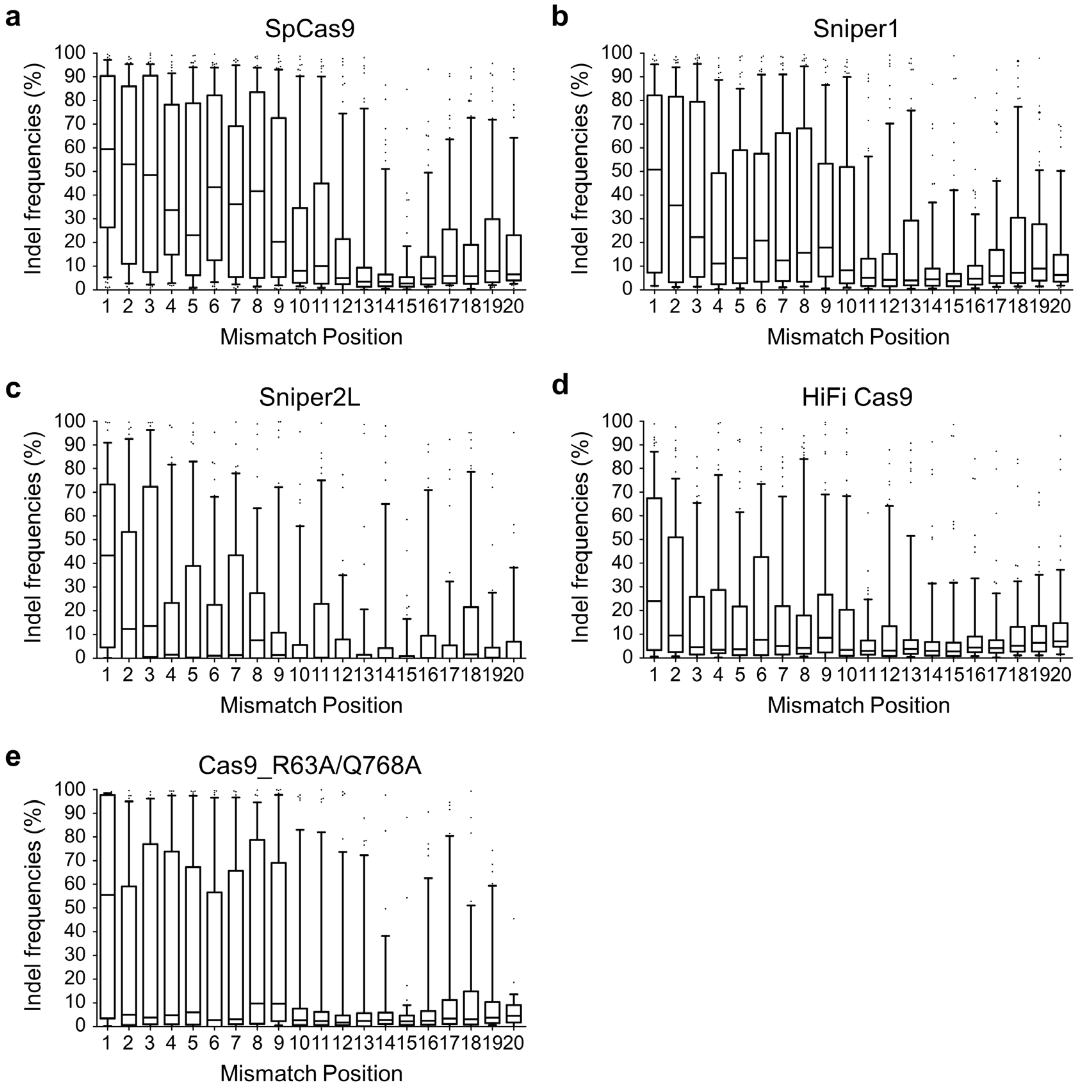


**Supplementary Figure 17.** Indel frequencies induced by high-fidelity Cas9 variants at target sequences containing single-base mismatches vary depending on the mismatch position. The boxes represent the 25th, 50th, and 75th percentiles; whiskers show the 10th and 90th percentiles. The number of target sequence *n* = 66 (position 1), 66 (position 2), 71 (position 3), 65 (position 4), 69 (position 5), 66 (position 6), 65 (position 7), 67 (position 8), 72 (position 9), 70 (position 10), 68 (position 11), 76 (position 12), 66 (position 13), 72 (position 14), 76 (position 15), 67 (position 16), 70 (position 17), 70 (position 18), 70 (position 19), and 62 (position 20) for SpCas9, 71 (position 1), 73, 75, 71, 76, 67, 70, 71, 73, 75, 77, 78, 71, 65, 80, 75, 71, 76, 74, and 62 (position 20) for Sniper0, 49 (position1), 41, 51, 44, 58, 52, 50, 47, 55, 53, 56, 48, 49, 48, 62, 49, 57, 64, 45, and 45 (position 20) for Sniper2L, 69 (position 1), 72, 68, 67, 72, 70, 68, 70, 76, 64, 67, 69, 67, 70, 78, 67, 75, 69, 71, and 64 (position 20) for HiFi Cas9, and 41 (position 1), 44, 41, 48, 45, 45, 42, 38, 48, 49, 46, 44, 43, 39, 49, 47, 47, 42, 45, and 32 (position 20) for Cas9_R63A/Q768A.

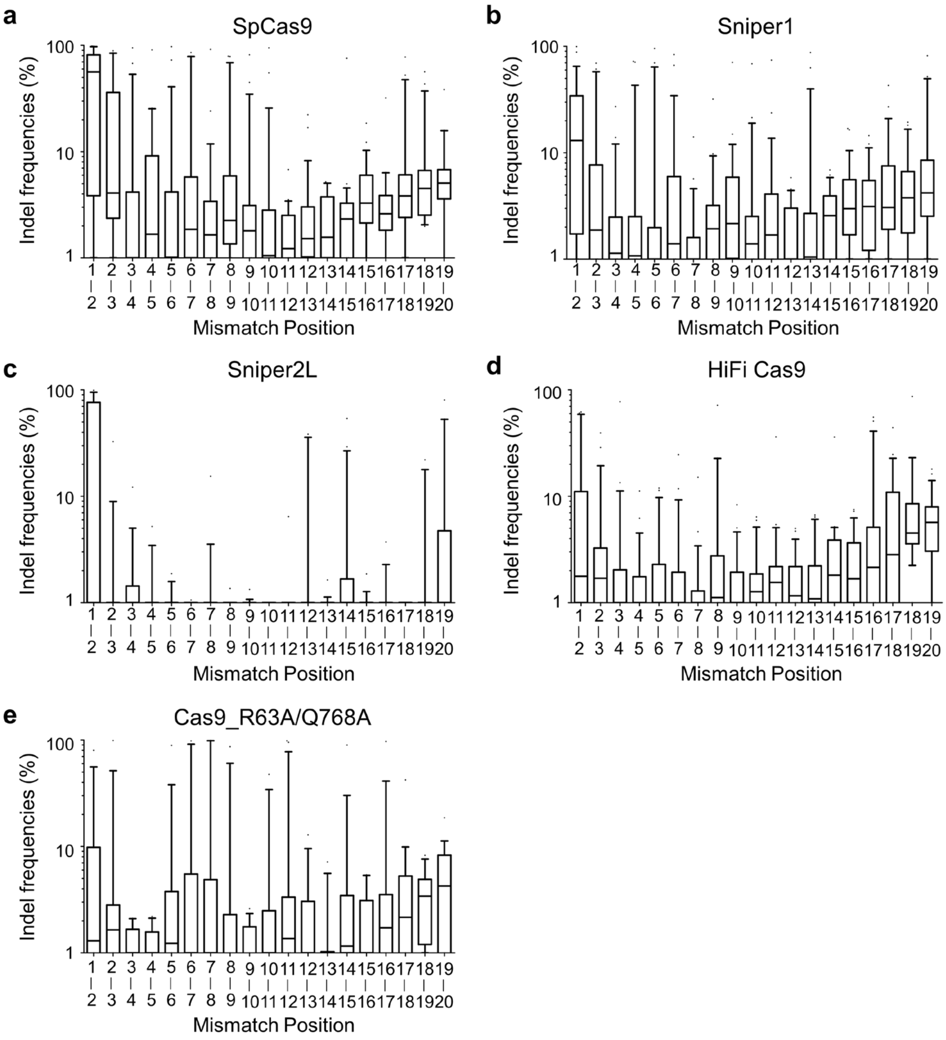


**Supplementary Figure 18.** Indel frequencies induced by high-fidelity Cas9 variants at target sequences containing consecutive two-base mismatches vary depending on the mismatch position. The boxes represent the 25th, 50th, and 75th percentiles; whiskers show the 10th and 90th percentiles. The number of target sequence *n* = 22 (position 1-2), 28 (position 2-3), 23 (position 3-4), 22 (position 4-5), 24 (position 5-6), 26 (position 6-7), 26 (position 7-8), 22 (position 8-9), 26 (position 9-10), 27 (position 10-11), 21 (position 11-12), 26 (position 12-13), 20 (position 13-14), 24 (position 14-15), 23 (position 15-16), 21 (position 16-17), 25 (position 17-18), 21 (position 18-19), and 21 (position 19-20) for SpCas9, 27 (position 1-2), 27, 21, 25, 27, 26, 25, 24, 24, 25, 25, 23, 23, 22, 25, 26, 23, 22, and 26 (position 19-20) for Sniper1, 20 (position 1-2), 18, 18, 18, 15, 18, 18, 13, 16, 17, 19, 13, 12, 20, 19, 16, 20, 11, and 21 (position 19-20) for Sniper2L, 29 (position 1-2), 24, 24, 25, 25, 22, 24, 23, 21, 24, 22, 24, 20, 20, 23, 23, 25, 19, and 24 (position 19-20) for HiFi Cas9, and 17 (position 1-2), 14, 13, 16, 15, 14, 14, 13, 13, 12, 21, 14, 12, 16, 11, 15, 19, 11, and 18 (position 19-20) for Cas9_R63A/Q768A.

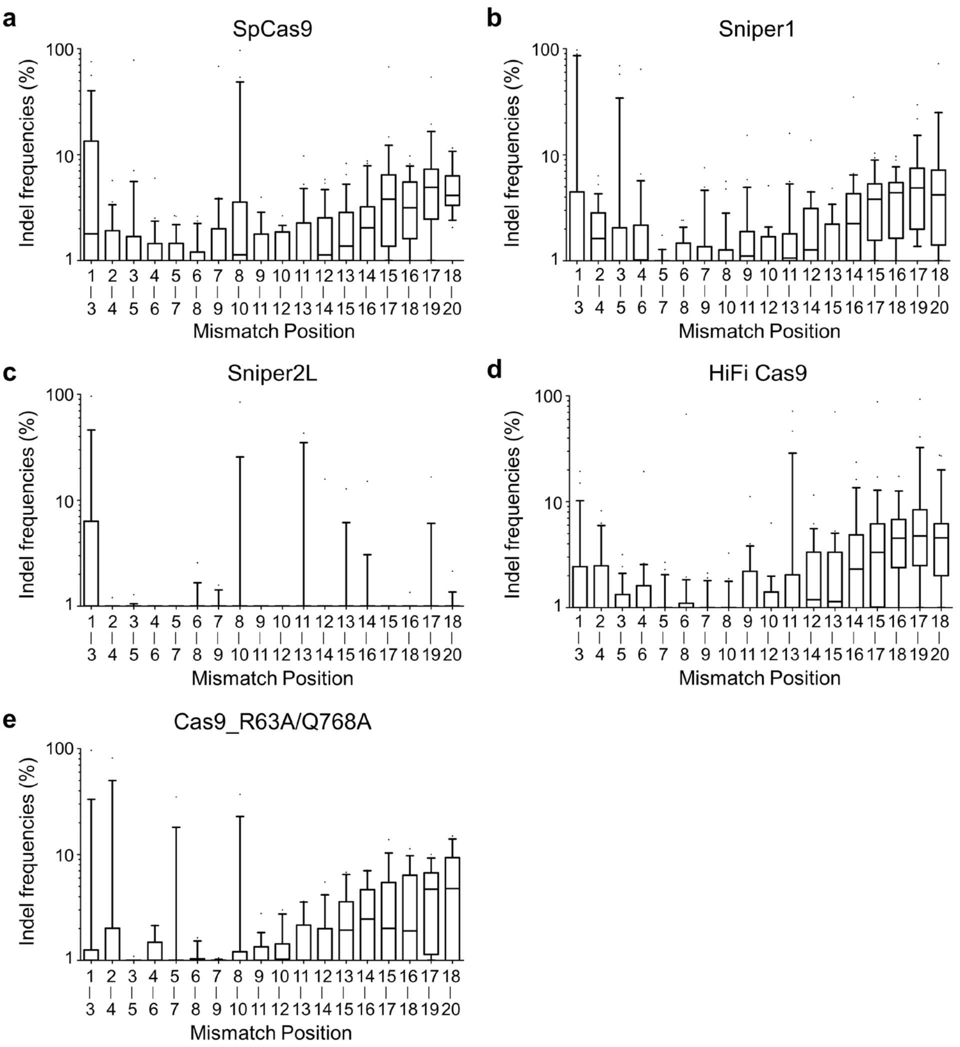


**Supplementary Figure 19.** Indel frequencies induced by high-fidelity Cas9 variants at target sequences containing consecutive three-base mismatches vary depending on the mismatch position. The boxes represent the 25th, 50th, and 75th percentiles; whiskers show the 10th and 90th percentiles. The number of target sequence *n* = 25 (position 1-3), 22 (position 2-4), 22 (position 3-5), 24 (position 4-6), 28 (position 5-7), 22 (position 6-8), 22 (position 7-9), 22 (position 8-10), 21 (position 9-11), 24 (position 10-12), 22 (position 11-13), 26 (position 12-14), 25 (position 13-15), 21 (position 14-16), 24 (position 15-17), 21 (position 16-18), 22 (position 17-19), and 21 (position 18-20) for SpCas9, 23 (position 1-3), 24, 25, 25, 27, 24, 22, 26, 23, 23, 20, 23, 25, 25, 26, 25, 29, and 20 (position 18-20) for Sniper1, 18 (position 1-3), 20, 12, 19, 18, 13, 10, 16, 14, 20, 12, 19, 18, 18, 21, 16, 17, and 15 (position 18-20) for Sniper2L, 24 (position 1-3), 21, 25, 23, 25, 23, 25, 21, 24, 20, 23, 24, 22, 22, 25, 19, 25, and 23 (position 18-20) for HiFi Cas9, and 16 (position 1-3), 13, 16, 11, 14, 11, 14, 13, 17, 11, 16, 15, 10, 10, 14, 16, 17, and 11 (position 18-20) for Cas9_R63A/Q768A.

**Supplementary Table 1.** Target sequences used in Sniper-screen (Provided as a separate excel file).

**Supplementary Table 2.** Indel frequencies at targets in library A when delivered as lentivirus (Provided as a separate excel file).

**Supplementary Table 3.** Indel frequencies at targets in library B when delivered as lentivirus (Provided as a separate excel file).

**Supplementary Table 4.** Indel frequencies at targets in library C when delivered as lentivirus (Provided as a separate excel file).

**Supplementary Table 5.** Indel frequencies at targets in library A when delivered as RNP (Provided as a separate excel file).

**Supplementary Table 6.** Primers used in this study (Provided as a separate excel file).

**Supplementary Table 7.** The sequence of DNA targets used in the smFRET assay (Provided as a separate excel file).
